# Exploiting HLA-II Promiscuity via Peptide Terminal Overhang Recognition for Pan-Allelic and Tumor-Selective AML Immunotherapy

**DOI:** 10.64898/2026.03.01.708866

**Authors:** Saori Fukao, Evey Y. F. Zheng, Fumie Ihara, Yukiko Matsunaga, Yota Ohashi, Dong-Hoon Han, Xinyu Wei, Kana Hasegawa, Brian D. Burt, Kayoko Saso, Dalam Ly, Marcus O. Butler, Mark D. Minden, Yuki Kagoya, Naoto Hirano

**Affiliations:** Princess Margaret Cancer Centre, University Health Network, Toronto, Ontario, Canada; Department of Immunology, University of Toronto, Toronto, Ontario, Canada; Department of Medicine, University of Toronto, Toronto, Ontario, Canada; Division of Tumor Immunology, Institute for Advanced Medical Research, Keio University, Tokyo, Japan; Centre for Oncology and Immunology, The University of Hong Kong, Hong Kong SAR, China

## Abstract

Antibodies targeting peptides presented by human leukocyte antigen (HLA) molecules expand the therapeutic landscape by enabling recognition of intracellular antigens. While most efforts have focused on allele-restricted peptides presented by HLA class I (HLA-I), HLA class II (HLA-II) epitopes remain underexplored despite their potential for promiscuous presentation. Acute myeloid leukemia (AML) is characterized by high expression of both HLA-II and the myeloid lineage antigen myeloperoxidase (MPO). Here, we identified an MPO-derived epitope (MPO_100-132_) that is promiscuously presented by multiple HLA-II molecules. We generated a MPO_100-132_-specific antibody (146D5) that recognizes the N-terminal overhang of this peptide independent of specific HLA contacts, enabling pan-allelic recognition. Engineered into bispecific T cell engagers (BiTEs), this antibody mediated robust cytotoxicity against primary AML samples across diverse HLA-II backgrounds. Crucially, 146D5-based BiTEs selectively spared normal myeloid cells, indicating that the MPO_100-132_ peptide, derived from the MPO propeptide, was functionally undetectable in normal myeloid cells, providing a significant safety window. *In vivo*, the MPO-targeting BiTE demonstrated potent antitumor activity and prolonged survival in AML xenograft models. Our findings identify peptide terminal overhangs as an actionable class of antibody targets and introduce a strategy to exploit HLA-II promiscuity for broadly applicable HLA-dependent but allele-agnostic immunotherapies.

## Introduction

Over the past century, antibody (Ab)-based therapies have been developed to treat a wide range of cancers^1,2^. Among these, T cell retargeting therapies, including T cell engagers (TCEs), chimeric antigen receptor (CAR)-T cells, have emerged as powerful modalities in cancer immunotherapy^3–7^. These approaches redirect cytotoxic T cells toward a specific surface antigen on tumor cells and promote the elimination of targeted cells. More recently, antibodies specific for tumor-derived peptides presented by major histocompatibility complex (MHC) proteins, known as T cell receptor (TCR) mimicking (TCRm) Abs, have further expanded the spectrum of targetable antigens to include intracellular proteins^8,9^.

TCRm Ab-based therapies have largely focused on peptides presented by MHC class I (MHC-I), yet their recognition is typically HLA-restricted, limiting broad applicability. This restriction reflects the structural features of MHC-I, which presents short peptides of 8-12 amino acids within a closed peptide-binding groove that anchors both termini and imposes strict allele-specific constraints on peptide presentation^10^. In addition, the binding modes of conventional TCRm Abs often reinforce HLA dependence. Some engage peptide–HLA (pHLA) complexes in a TCR-like manner, whereas others predominantly contact the HLA scaffold rather than the peptide itself^11,12^. Although a limited number of studies have reported Abs or CAR derivatives capable of recognizing a peptide presented across multiple HLA-I alleles^12,13^, such cross-recognition is generally limited to closely related allelic subtypes with similar interface topologies^12,14^. In principle, Abs can engage diverse epitopes within pHLA complexes with high affinity. Accordingly, strategies specifically designed to drive peptide-centric recognition independent of HLA subtype may overcome HLA restriction and expand the clinical applicability of pHLA-targeted immunotherapies.

Unlike MHC-I, MHC class II (MHC-II) molecules have an open-ended peptide-binding cleft that accommodates longer peptides, typically 13–25 amino acids, with N- and C-terminal overhangs or even larger unfolded proteins via peptide-like regions^10,15^. This structural feature allows promiscuous peptide presentation, enabling a single peptide to be presented by multiple HLA-II alleles via alternative binding registers^16–18^. For example, the class II-associated invariant chain peptide (CLIP) interacts with diverse HLA-II molecules through different binding modes^19–21^. Immunopeptidomic analyses have further shown that certain peptides are promiscuously presented across structurally different HLA-II groups^18^. Such peptides may represent attractive targets for Ab-based therapies aimed at circumventing HLA restrictions. Although HLA-II is predominantly expressed on professional antigen-presenting cells (APCs), growing evidence indicates that HLA-II is also expressed on many tumor types, including hematological malignancies and solid tumors^22,23^, and can present endogenous antigens processed through alternative pathways^24–27^. TCR gene therapies targeting HLA-II presented peptides have demonstrated clinical efficacy^28,29^, further underscoring the therapeutic potential of HLA-II-restricted antigens.

Acute Myelogenous Leukemia (AML), one of the most common leukemias in adults, frequently enters remission but commonly recurs and remains highly lethal. AML cells often exhibit high expression of both HLA-II and MPO, a hallmark enzyme of myeloid lineage commitment^30–32^. Although the therapeutic potential of MPO in AML has been explored in the context of HLA-I ^33^, MPO-derived peptides presented by HLA-II remain largely underexplored as therapeutic targets for AML. Here, we have identified a novel MPO-derived epitope that is promiscuously presented across multiple HLA-II alleles and developed an Ab-based bispecific TCE (BiTE) capable of recognizing this epitope in an HLA-dependent but allele-agnostic manner to treat a broad spectrum of AML patients.

## Results

### Identification of an MPO peptide promiscuously presented across multiple HLA-II alleles to target a broad range of AML samples

In AML, expression of HLA-II, particularly HLA-DR, together with MPO is an important diagnostic marker and is used for disease classification^30,31,34^. Transcriptomic analyses of large AML patient cohorts^35,36^ revealed frequent co-expression of HLA-II and MPO at high levels (Fig. 1a). To validate protein expression, we analyzed primary AML patient samples by flow cytometry. Consistent with previous reports^30,31^, most MPO^+^ AML blasts exhibited high HLA-II expression (Fig. 1b). Thus, MPO is abundantly expressed in HLA-II^+^ AML cells across many patients, suggesting that MPO-derived peptides could be presented by HLA-II and may represent promising targets for T cell-based therapies.

**Fig. 1.**
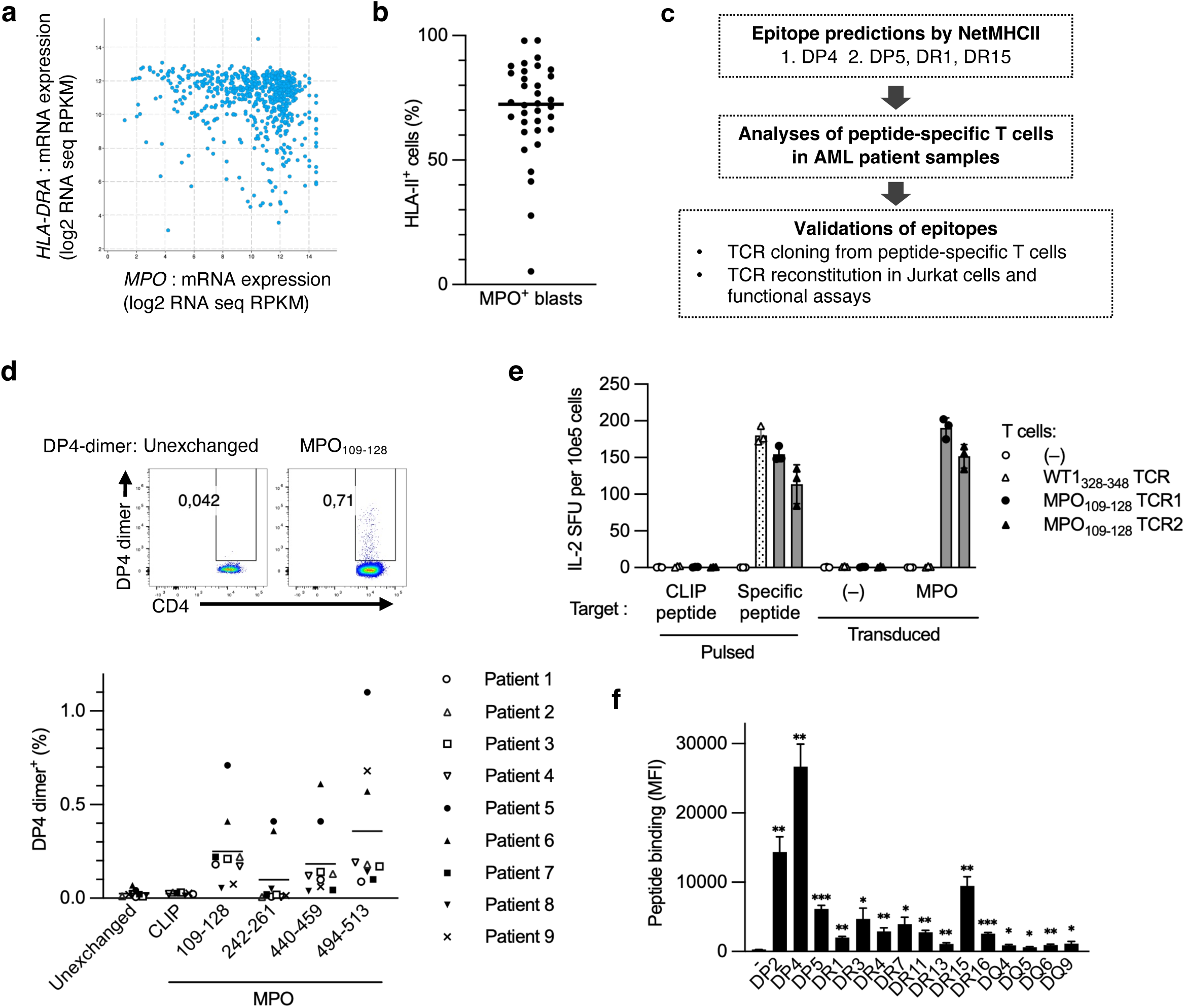
MPO peptides presented by HLA-DP4 as a potential immunotherapeutic target in AML. a,mRNA expression of *MPO* and *HLA-DRA* analyzed using cBioPortal for Cancer Genomics database. **b,** Frequency of HLA-II^+^ cells among MPO^+^ blasts in AML patient samples analyzed by flow cytometry. Bar indicates mean. **c,** Schematic of the discovery and validation workflow for a promiscuously presented MPO-derived peptide epitope by HLA-DP4 and other HLA-II alleles. **d,** Representative staining (top) and frequencies (bottom) of CD4^+^ T cells specific for each MPO-derived peptide/DP4 dimer from DP4^+^ AML patients. **e,** Jurkat76/CD4 cells transduced with DP4-restricted TCRs specific for WT1_328-348_ or MPO_109-128_ (cloned from DP4^+^ donors) or untransduced controls were cocultured with DP4^+^ K562 cells pulsed with CLIP or cognate peptides, or with DP4^+^ K562 cells transduced with full-length MPO or untransduced. IL-2 spot-forming units (SFU) per 10^5^ cells are shown. **f,** Binding of biotinylated MPO_100-132_ peptide to K562 cells expressing the indicated HLA-II alleles, detected by streptavidin staining (MFI). P-values were calculated by one-way ANOVA with Dunnett’s multiple comparisons, using HLA-II negative K562 cells (–) as the control group: ***p<0.001, **p<0.01, *p<0.05. Data represent mean from 9 donors (d) or mean ± S.D. from 3 (e) or 4 (f) technical replicates. Pooled data from 2 independent experiments (b,d) or representative data from 2 independent experiments (e,f).

To explore whether MPO-derived peptides can bind HLA-II, we performed *in silico* peptide binding predictions using NetMHCII^37,38^. Since HLA-DPA1*01:03/DPB1*04:01 (DP4) is the most frequent allele globally^39,40^, we initially focused on peptides predicted to bind DP4. Additional filters identified peptides predicted to bind other common HLA-II alleles, including HLA-DR variants, thereby enriching for candidates with potential promiscuous binding across multiple HLA-II alleles (Fig. 1c, Supplementary Table 2). Based on predicted binding affinity (half-maximal inhibitory concentration, IC_50_ < 500 nM), four peptides were selected for further analysis.

We then tested whether MPO peptides are endogenously processed and presented by HLA-II and recognized by T cells in AML patients. Using peptide-loaded DP4 dimers^41^, we detected CD4^+^ T cells specific for each peptide at varying frequencies among DP4^+^ AML patient samples (Fig. 1d). To confirm TCR specificity toward endogenously presented peptides, we cloned TCRs from DP4 dimer-positive T cells and reconstituted them in Jurkat cells. Jurkat cells expressing an MPO_109-128_-specific TCR responded to K562 cells co-expressing MPO and DP4, but not to MPO^-^ cells (Fig. 1e). These results indicate that MPO_109-128_ is endogenously processed and presented by DP4 in MPO^+^ tumor cells and can be recognized by T cells.

We next determined whether the extended peptide MPO_100–132_ (encompassing MPO_109-128_) could bind multiple HLA-II alleles. Biotinylated MPO_100-132_ exhibited strong binding to DP alleles, moderate binding to several DR alleles, and weak binding to DQ alleles expressed in K562 cells (Fig. 1f). *In silico* predictions supported these findings, showing that MPO_100-132_ engages distinct HLA-II alleles through different binding cores with varying affinities (Supplementary Table 3). These results demonstrate that MPO_100-132_ can bind promiscuously to multiple HLA-II alleles.

### Development of a peptide-specific antibody targeting the MPO peptide presented by HLA-DP4

To design an antibody with broad HLA recognition, we aimed to generate one that recognizes the peptide itself, independent of HLA contacts. HLA-II-bound peptides typically extend beyond the binding cleft at both termini, exposing N- and C-terminal overhangs that may be accessible for antibody recognition^10^. These overhanging residues are less structurally constrained by the HLA-II interaction, making them promising targets for antibodies capable of recognizing the same peptide across multiple HLA-II alleles. To generate an antibody that could recognize the MPO peptide overhang independently of HLA contacts, we immunized mice with the N- (MPO_100-112_) and C-terminal (MPO_122-132_) overhang peptides derived from MPO_100-132_, and generated hybridomas (Fig. 2a, Extended Data Fig. 1). Among the screened hybridomas, 22 recognized MPO_100-132_ in ELISA analysis, 8 recognized MPO_100-132_ pulsed on DP4^+^ K562 cells, and 3 recognized MPO peptide endogenously presented by K562 cells co-expressing MPO and DP4 (Extended Data Fig. 1). Clone 146D5 exhibited consistent binding across all tested conditions and was selected for further characterization. The 146D5 antibody (146D5 Ab) specifically bound to MPO_100-132_ as well as peptide-pulsed on DP4^+^ K562 cells (Fig. 2b,c). Moreover, it bound K562 cells co-expressing DP4 and MPO, but not cells lacking either molecule (Fig. 2d), demonstrating recognition of endogenously processed and presented peptide. Bio-layer interferometry (BLI) revealed an affinity of 146D5 Ab for the MPO_100–132_–DP4 complex of ∼25 nM (Fig. 2e), which is within the range of other therapeutic TCRm Abs^13,42^.

**Fig. 2.**
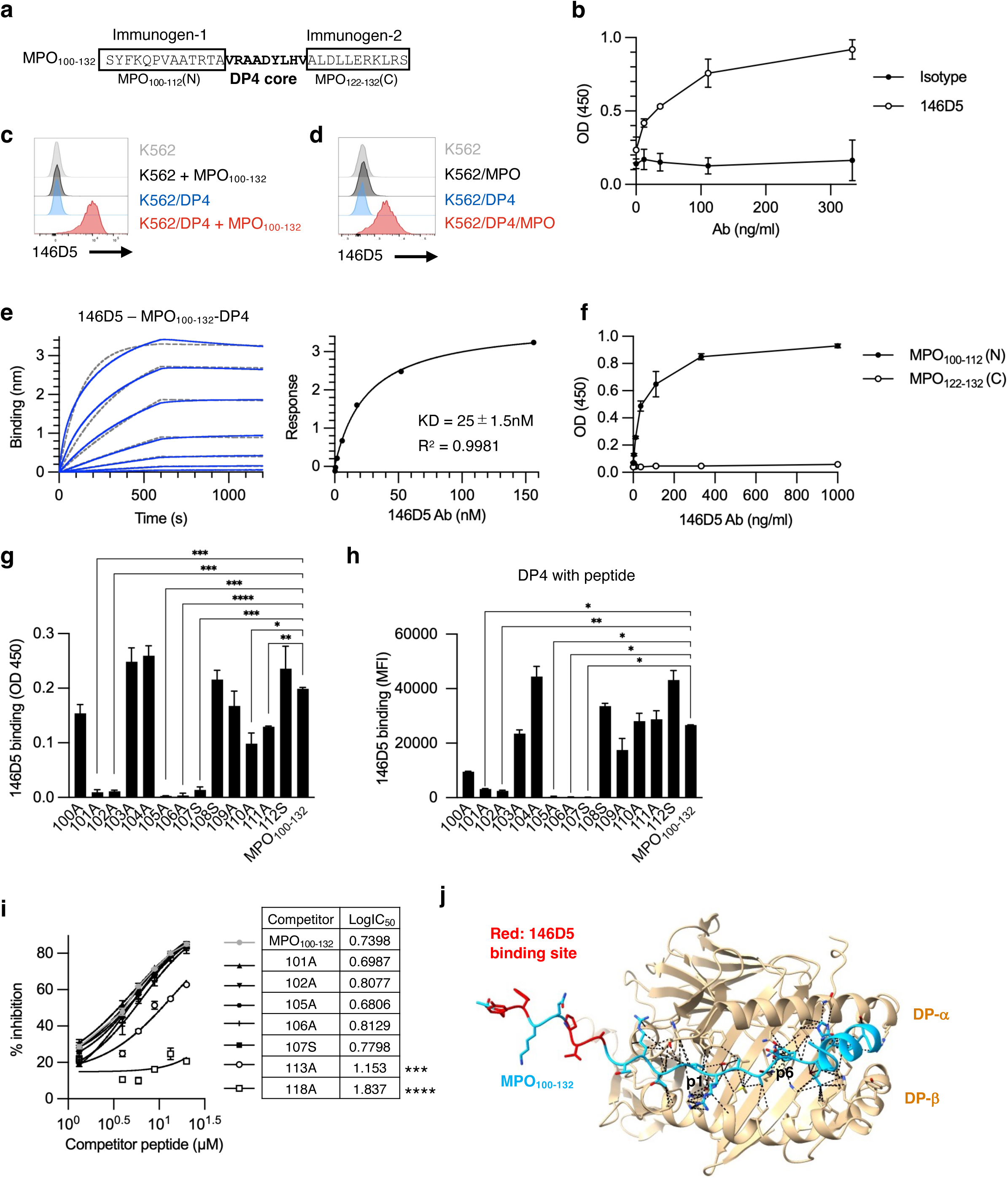
Generation and characterization of 146D5 Ab specific for MPO_100-132_ a, Amino acid sequence of the MPO_100-132_ peptide with predicted DP4 binding core (bold) and immunogen regions (boxed). **b,** Binding of 146D5 Ab to plate-immobilized MPO_100-132_ via ELISA. **c,d,** Binding of 146D5 Ab to K562 cells with or without DP4 expression, pulsed with MPO_100-132_ (c) or K562 cells untransduced or transduced with DP4, MPO, or both (d). **e,** Representative BLItz sensorgram showing the binding of 146D5 Ab to biotinylated MPO_100-132_ / DP4 complex. **f,g,** ELISA analysis of 146D5 Ab binding to N- or C-terminal MPO-derived peptides (f) and alanine/serine-substituted variants of MPO_100-132_ (g). **h,** Binding of 146D5 Ab to DP4^+^ K562 cells pulsed with alanine/serine-substituted MPO_100-132_ mutants. P-values by one-way ANOVA followed by Dunnett’s multiple comparisons (g,h): ****p<0.0001, ***p<0.001, **p<0.01, *p<0.05. **i**, Competitive binding assays using DP4^+^ K562 cells pulsed with graded concentrations of MPO_100-132_ or its mutants. LogIC_50_ values were calculated by nonlinear regression with variable slope fitting. LogIC_50_ values from replicate measurements were compared by one-way ANOVA followed by Dunnett’s multiple comparisons, using MPO_100-132_ as the control group. ****p<0.0001, ***p<0.001. **j,** Structural model of MPO_100-132_ bound to DP4 generated using AlphaFold. The peptide is shown in cyan with 146D5 Ab-contact residues in red; HLA-II in light brown. Data represent mean ± S.D. from technical duplicates (b, f-i). Representative data from at least 2 independent experiments are shown (b-i).

### Identification of the 146D5 Ab epitope in the MPO peptide-DP4 complex

To define the antibody epitope, we analyzed 146D5 binding to the MPO_100-132_ N- and C-terminal overhang regions. 146D5 Ab specifically recognized the N-terminal peptide but not the C-terminal one (Fig. 2f). Alanine (or serine, when the original residue was alanine) scanning revealed that substitutions at Y101, F102, P105, V106, and A107 of the MPO peptide markedly decreased 146D5 Ab binding to both peptide and peptide-pulsed DP4^+^ K562 cells, whereas substitutions at adjacent residues modestly reduced binding (Fig. 2g,h). These findings indicate that these five residues are critical for recognition, with neighboring residues providing secondary contributions.

To confirm that reduced binding was not due to impaired peptide loading onto DP4, we performed competitive binding assays. Peptide variants carrying alanine/serine substitutions at Y101, F102, P105, V106, and A107 bound DP4-expressing cells at comparable levels to wild-type peptide, as shown by LogIC_50_ values (Fig. 2i). In contrast, substitutions at V113 and Y118, predicted p1 and p6 anchors within the DP4 core (Supplementary Table 3)^39^, substantially impaired binding to DP4 (Fig. 2i).

Structural modeling of MPO_100-132_ bound to DP4 using AlphaFold^43^ supported these observations: V113 and Y118 engaged with DP4 cleft, while Y101, F102, P105, V106, and A107 formed an exposed N-terminal overhang accessible to 146D5 (Fig. 2j, Extended Data Fig. 2a). Together, these results indicate that 146D5 Ab recognizes a distinct N-terminal overhang epitope of MPO_100-132_ presented by DP4, independent of HLA contacts.

### 146D5 Ab cross-recognizes MPO peptide presented by multiple HLA-II

We next evaluated whether 146D5 Ab could recognize MPO_100-132_ presented by other HLA-II alleles. Consistent with the peptide binding profile observed in HLA-II expressing K562 cells (Fig. 1f), 146D5 Ab strongly bound MPO_100-132_ pulsed on DP alleles and showed moderate binding to DR alleles (Fig. 3a). To confirm recognition of endogenously presented peptide, we analyze 146D5 binding to K562 cells co-expressing MPO with various HLA-II alleles (Fig. 3b). Significant binding was again detected across multiple HLA-II alleles. For certain DR and DQ alleles, endogenously processed MPO elicited stronger binding of 146D5 Ab than exogenous peptide pulsing (Fig. 3a,b). This may reflect allele-specific differences in antigen processing, whereby naturally generated MPO-derived peptides could vary in register, length, or post-translational modification relative to the synthetic MPO_100–132_ peptide used for pulsing, potentially altering epitope display and antibody recognition. Further validation was obtained by staining of K562 cells expressing 146D5-IgG with MPO_100–132_-loaded HLA-II tetramers (Fig. 3c). Tetramers for DP4, DR1, and DR15 bound specifically to 146D5-IgG–expressing cells but not to control cells (Fig. 3c).

**Fig. 3.**
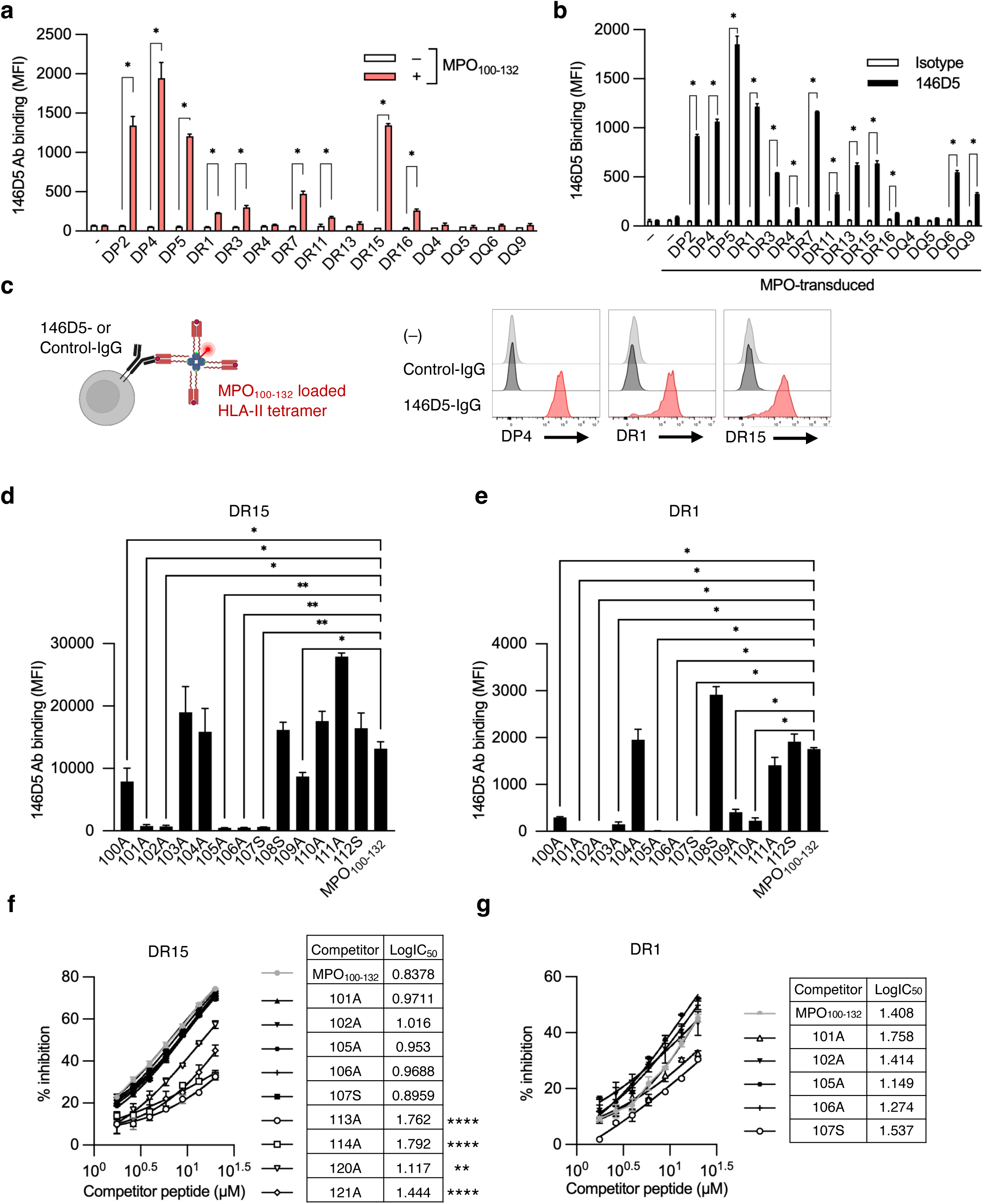
146D5 Ab recognizes MPO peptide presented by multiple HLA-II alleles **a-b,** Binding of 146D5 Ab to K562 cells expressing the indicated HLA-II alleles and pulsed with MPO_100-132_ (a) or transduced with MPO (b). Data represent mean ± S.D. from technical duplicates. P-values were determined by Multiple unpaired t tests: *p<0.05. **c,** Schematic (left) and binding of MPO_100-132_ loaded DP4, DR1, or DR15 tetramers to K562 cells expressing control-IgG or 146D5-IgG (right). **d,e,** Binding of 146D5 Ab to DR15^+^ (d) or DR1^+^ (e) K562 cells pulsed with MPO_100-132_ mutants. Data represent mean ± S.D. from technical duplicates. P-values were determined by one-way ANOVA followed by Dunnett’s multiple comparisons: **p<0.01, *p<0.05. **f,g,** Competitive binding assays using DR15^+^ (f) or DR1^+^ (g) K562 cells pulsed with graded concentrations of MPO_100-132_ or mutants. LogIC_50_ values from nonlinear regression. LogIC_50_ values from replicate measurements were compared by one-way ANOVA followed by Dunnett’s multiple comparisons, using MPO_100-132_ as the control group. ****p<0.0001, **p<0.01. Representative of at least 2 independent experiments (a-g).

Epitope mapping in the context of DR1 and DR15 revealed that substitutions at Y101, F102, P105, V106, and A107 again impaired 146D5 Ab binding (Fig. 3d,e). In DR15, substitutions at the anchor residues p1, p2, p8, and p9—corresponding to V113, R114, H120, and V121—reduced peptide binding to HLA molecules, whereas substitutions at other residues did not^45,46^ (Fig. 3f). This is consistent with AlphaFold modeling showing that these residues reside within the DR15 binding cleft, whereas the 146D5 epitope is exposed on the N-terminal overhang (Extended Data Fig. 2b,c).^46,47^ In DR1, overlap between the 146D5 epitope and the predicted DR1 binding core were suggested (Extended Data Fig. 2d). Substitutions at these five residues seemed to affect peptide binding to HLA molecules, although not significantly (Fig. 3g). MPO_100-132_ bound more weakly to DR1 than to DP4 and DR15 (Fig. 1f), which may have limited detection of differences. Partial burial of the peptide within the DR1 binding groove or alternative binding modes may permit access of 146D5 to the epitopes. Overall, MPO_100-132_ is presented by various HLA-II alleles through distinct anchor residues, while the N-terminal overhang remains accessible to 146D5 Ab, enabling broad cross-recognition.

### 146D5 BiTE induces T cell responses against MPO^+^ cells expressing diverse HLA-II molecules

To harness 146D5 for therapy, we engineered a BiTE by fusing the single-chain variable fragment (scFv) from 146D5 with the scFv from anti-CD3 (OKT3) (Fig. 4a). 146D5 BiTE bound to DP4^+^ K562 cells pulsed with MPO_100-132_, as well as to K562 cells co-expressing MPO and DP4 (Fig. 4b,c), and CD3^+^ T cells (Fig. 4d). In functional assays, 146D5 BiTE redirected polyclonal T cells towards DP4^+^ K562 cells pulsed with MPO_100-132_, as well as K562 cells co-expressing MPO and DP4, resulting in concentration-dependent IFN-ψ production and cytotoxicity (Fig. 4e,f). As previously shown for similar BiTE formats, as little as 1-10 picomolar 146D5 BiTE was sufficient to elicit significant T cell responses^4^. Importantly, 146D5 BiTE redirected T cells to kill MPO^+^ K562 cells co-expressing various DP, DR, and DQ molecules (Fig. 4g). For some alleles such as DR3 or DQ4, cytotoxicity appeared stronger than expected based on the binding profile of 146D5 Ab (Fig. 3b). This likely reflects the higher sensitivity of T-cell cytotoxicity driven by BiTE engagement compared with monomeric antibody binding.

**Fig. 4.**
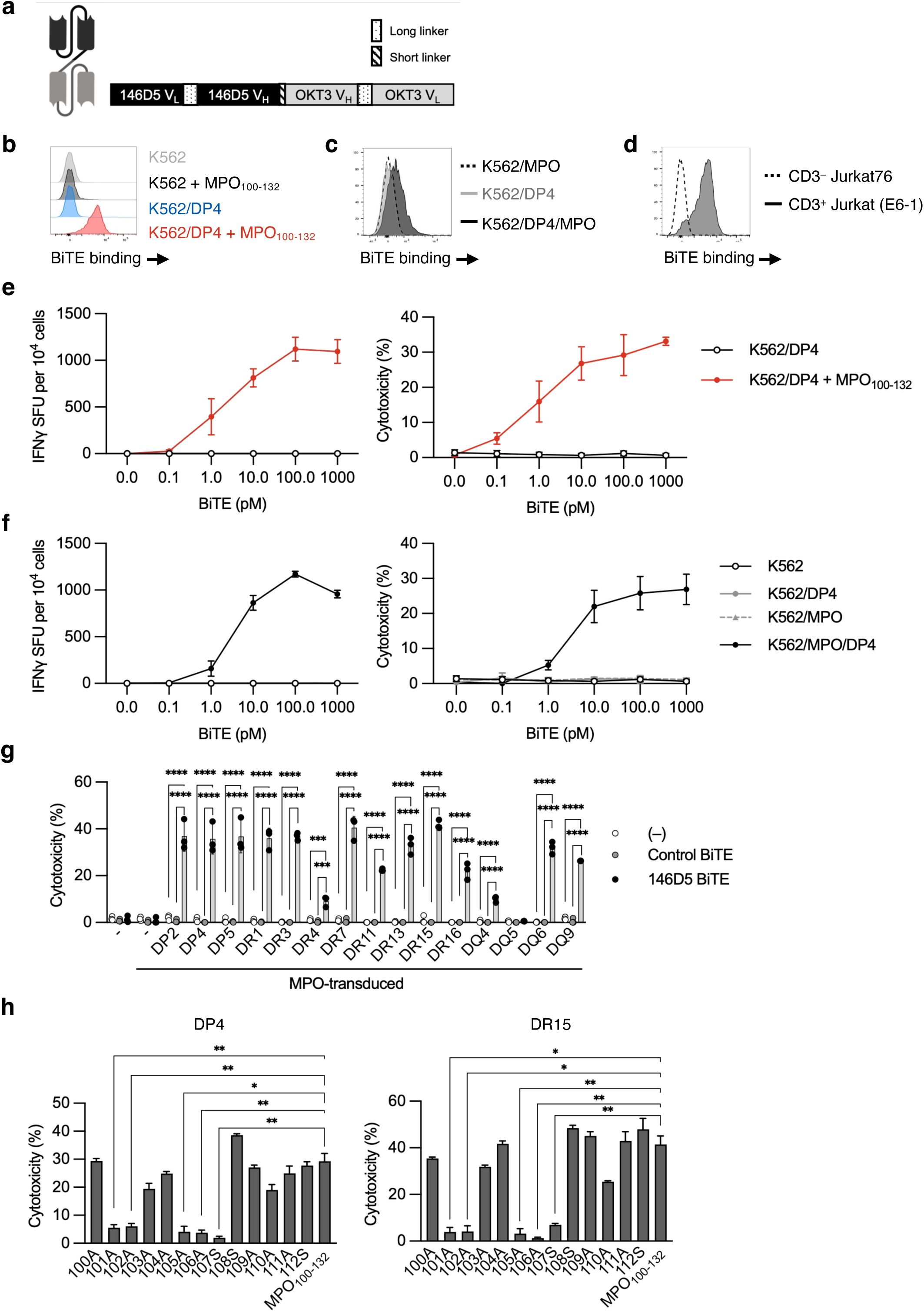
146D5 BiTE mediates T cell cytotoxicity against MPO^+^ targets across diverse HLA-II alleles a, Schematic of 146D5 BiTE. **b-d,** Binding of 146D5 BiTE to K562 cells with or without DP4 expression pulsed with MPO_100-132_ (b), K562 cells transduced with DP4, MPO, or both (c), CD3^+^ Jurkat (E6-1) cells, or CD3^-^ Jurkat76 cells (d). **e,f,** IFNψ SFU per 10^4^ cells (left) and cytotoxicity (right) induced by T cells cocultured with DP4^+^ K562 cells pulsed with MPO_100-132_ (e), or K562 cells untransduced or transduced with DP4, MPO, or both (f) in the presence of graded concentrations of 146D5 BiTE. **g,** Cytotoxicity induced by T cells cocultured with K562 cells expressing MPO and indicated HLA-II alleles in the presence of 1 nM indicated BiTE. P-values by two-way ANOVA followed by Šídák’s multiple comparisons: ****p<0.0001, ***p<0.001. **h,** Cytotoxicity induced by T cells cocultured with K562 cells expressing DP4^+^or DR15^+^ K562 cells pulsed with MPO_100-132_ mutants in the presence of 1 nM 146D5 BiTE. P-values by one-way ANOVA followed by Dunnett’s multiple comparisons: ***p<0.001, **p<0.01, *p<0.05. All cytotoxicity assays at E:T=2:1. Data represent mean ± S.D. from T cells from 3 different donors (e-h). Representative data of 2 independent experiments.

As with the parental antibody, substitutions at Y101, F102, P105, V106, and A107 abolished cytotoxicity across DP4, DR1, DR15, and DQ6.1 (Fig. 4h, Extended Data Fig. 3c,d). Collectively, these results demonstrate that 146D5 BiTE retains the specificity of 146D5 Ab while effectively redirecting T cells to kill MPO^+^ cells across multiple HLA-II molecules.

### 146D5 BiTE induces T cell responses against diverse AML samples

We next examined the activity of 146D5 BiTE against various AML samples. 146D5 BiTE induced cytotoxicity against AML cell lines, ME-1, Kasumi-1 and Nomo1, which express both MPO and diverse HLA-II alleles (Fig. 5a, Extended Data Fig. 4a, Supplementary Table 4). It also killed primary AML patient samples with high frequencies (>60%) of MPO^+^ and HLA-II^+^ blasts (Fig. 5b, Extended Data Fig. 4b, Supplementary Table 5). While MPO expression was uniformly high across the tested samples (Supplementary Table 5), the extent of cytotoxicity correlated with HLA-II expression levels (Fig. 5c).

**Fig. 5.**
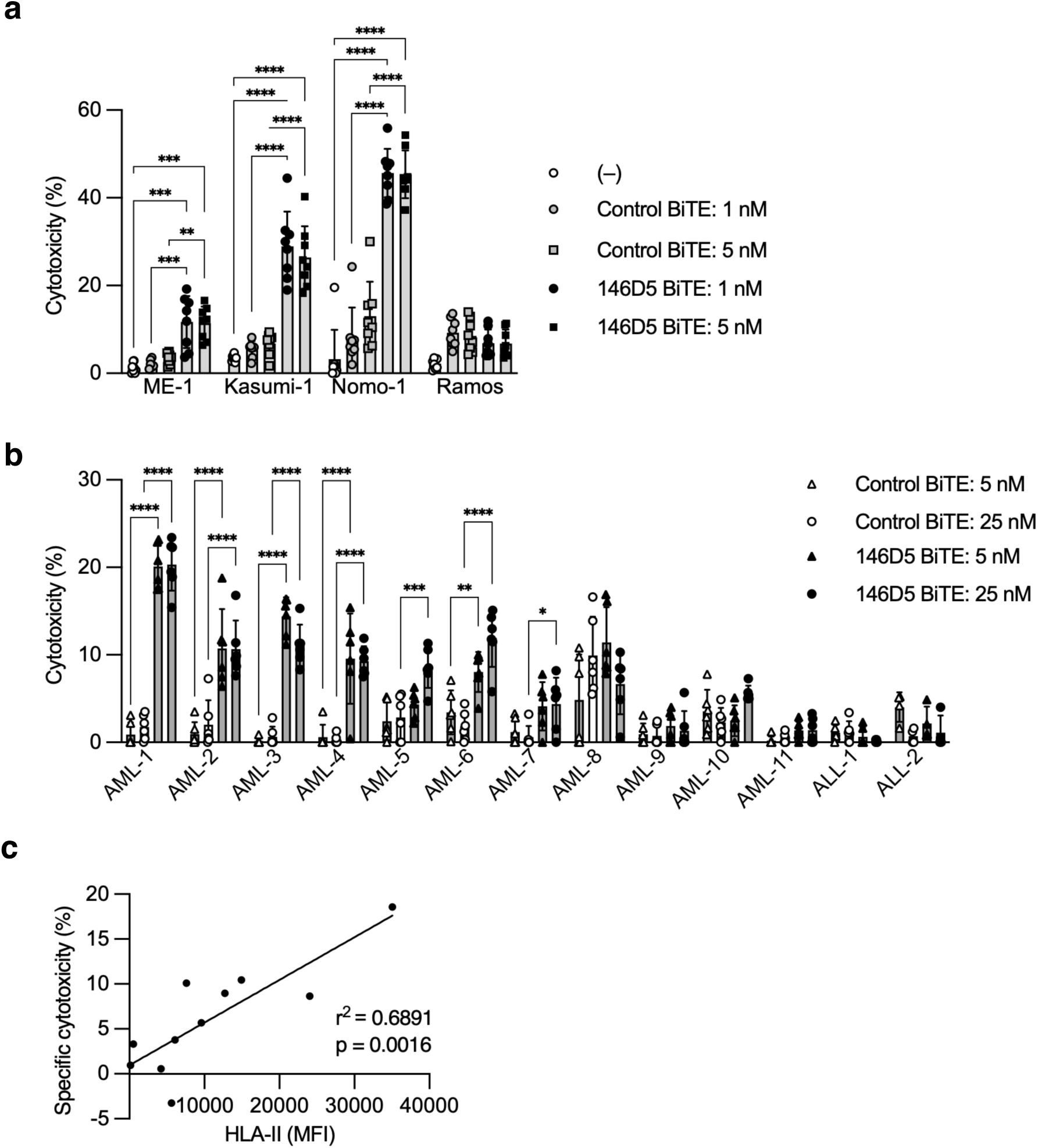
Recognition of diverse AML samples by 146D5 BiTE **a,b,** Cytotoxicity of cell lines at E:T=2:1 (a) or primary AML or ALL samples at E:T=4:1 (b) by T cells cocultured in the absence or presence of control BiTE or 146D5 BiTE. Data represent mean ± S.D. from T cells of 8 (a) or 6 (b) donors. P-values by two-way ANOVA followed by Turkey’s multiple comparisons: ****p<0.0001, ***p<0.001, **p<0.01, *p<0.05. **c,** Correlation between HLA-II expression (MFI) in AML samples and p-value by simple linear regression. Specific cytotoxicity (146D5 BiTE –control BiTE) using autologous T cells. Pooled data from 3 independent experiments (a-c).

Importantly, 146D5 BiTE did not kill MPO-negative acute lymphoblastic leukemia (ALL) samples (Fig. 5b), confirming target specificity.

### 146D5 BiTE lacks activity against normal myeloid cells

MPO is highly expressed in mature neutrophils and at lower levels in monocytes^44^. To assess potential off-target reactivity, we evaluated the activity of 146D5 BiTE against MPO-expressing normal myeloid cells. Naïve or activated neutrophils and monocytes were co-cultured with polyclonal T cells in the presence of BiTEs. As previously reported, high MPO expression was observed in naïve and activated neutrophils and in naïve monocytes, with lower levels in activated monocytes (Extended Data Fig. 4c,d). Although naïve neutrophils lacked constitutive HLA-II expression, activated neutrophils and monocytes expressed HLA-II (Extended Data Fig. 4c,d). Nevertheless, 146D5 BiTE did not induce cytotoxicity against any of these normal cells (Fig. 6a). In contrast, MPO_100–132_–pulsed monocytes were efficiently killed following 146D5 BiTE treatment, indicating insufficient endogenous peptide presentation in normal cells (Fig. 6b).

**Fig. 6.**
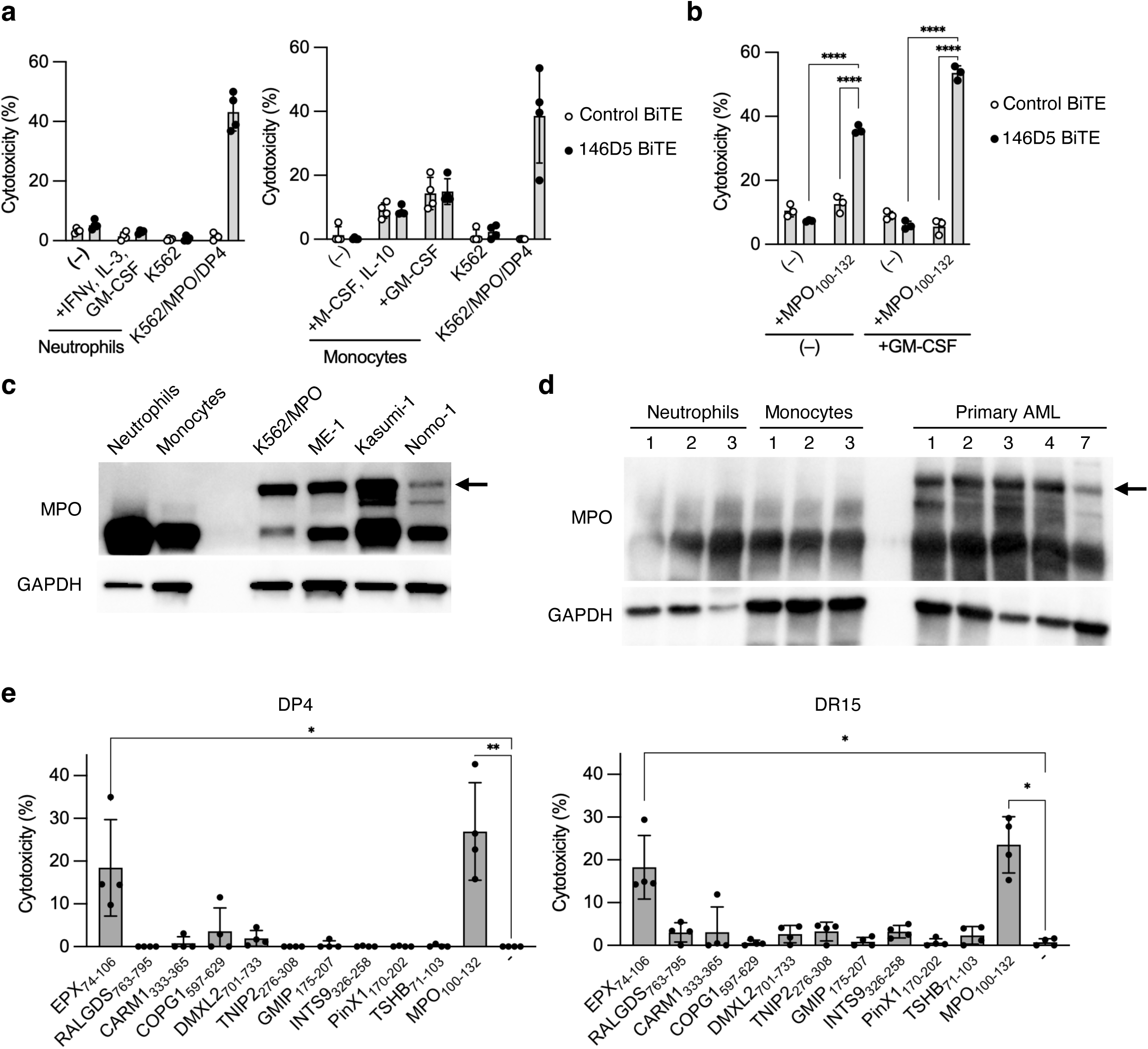
146D5 BiTE shows limited activity toward normal myeloid cells **a,b,** Cytotoxicity of naïve or activated neutrophils (a, left) or monocytes (a, right), or MPO_100-132_-pulsed monocytes (b) by T cells co-cultured with 5 nM BiTE. Neutrophils were stimulated for 3 days (a), and monocytes stimulated for 6 days (a) or 2 days (b) with indicated stimuli before assays. E:T=4:1. Cytotoxicity was shown. Data represent mean ± S.D. from 4 donors (a) or from 3 technical replicates (b). **c,d,** Immunoblots of MPO and GAPDH in cell lysates derived from neutrophils or monocytes from healthy donors, cell lines, and primary AML samples. Arrow indicates apoproMPO/proMPO. **e,** Cytotoxicity of DP4^+^ or DR15^+^ K562 cells pulsed with indicated peptides by T cells co-cultured in the presence of 1 nM 146D5 BiTE (E:T=2:1). Data represent mean ± S.D. from 4 donors. P-values by two-way ANOVA (b) or one-way ANOVA (e) followed by Šídák’s multiple comparisons: ****p<0.0001, **p<0.01, *p<0.05. Pooled data from 2 independent experiments (a, e) or representative data from at least 2 independent experiments (b-d) were shown.

MPO_100-132_ lies within the propeptide segment of MPO, which is normally removed during the processing of immature precursors such as apoproMPO and proMPO, into the subunits of mature MPO^8^. Because increased expression of immature precursors might enhance the presentation of MPO_100-132_, we compared MPO protein expression in 146D5 BiTE-susceptible AML cell lines and primary AML samples with that in non-susceptible normal myeloid cells. As expected, apoproMPO/proMPO was detected in AML cell lines and primary AML samples, but not in normal neutrophils or monocytes (Fig. 6c,d). These results suggest that normal myeloid cells fail to present MPO_100-132_ on HLA-II due to limited accumulation of immature MPO precursors. This difference may reflect more efficient protein processing in normal myeloid cells compared to AML.

We further assessed whether 146D5 BiTE exhibit cross-reactivity toward peptides that either share high sequence homology with the N-terminal overhang of MPO_100-132_ in the context of DP4, or contain the five key residues (Y101, F102, P105, V106, and A107) for 146D5 BiTE recognition (Fig. 4h). Among the tested homologous peptides (Supplementary Table 6), 146D5 BiTE redirected T cell cytotoxicity only towards eosinophil peroxidase (EPX)_74-106_ peptide-pulsed DP4^+^ or DR15^+^ K562 cells (Fig. 6e). EPX, a member of peroxidase superfamily, shares 70% overall sequence identity with MPO^45^, including 92% identity of corresponding region with N-terminal overhang of MPO_100-132_. Although EPX is expressed in eosinophils, they lack HLA-II expression (Extended Data Fig. 4e) and are unlikely to be recognized by 146D5 BiTE.

### 146D5 BiTE demonstrates significant efficacy in *in vivo* AML mouse models

Finally, we evaluated the *in vivo* therapeutic efficacy of 146D5 BiTE in two AML xenograft models. Kasumi-1 or Nomo-1 were transplanted into NSG mice, followed by the adoptive transfer of polyclonal T cells and BiTE treatment (Fig. 7a). Compared with untreated or control BiTE-treated groups, 146D5 BiTE treatment significantly suppressed tumor growth (Fig. 7b,d, Extended Data Fig. 5a,b) and prolonged mouse survival (Fig. 7c,e). These results demonstrate potent *in vivo* anti-leukemic efficacy of 146D5 BiTE.

**Fig. 7.**
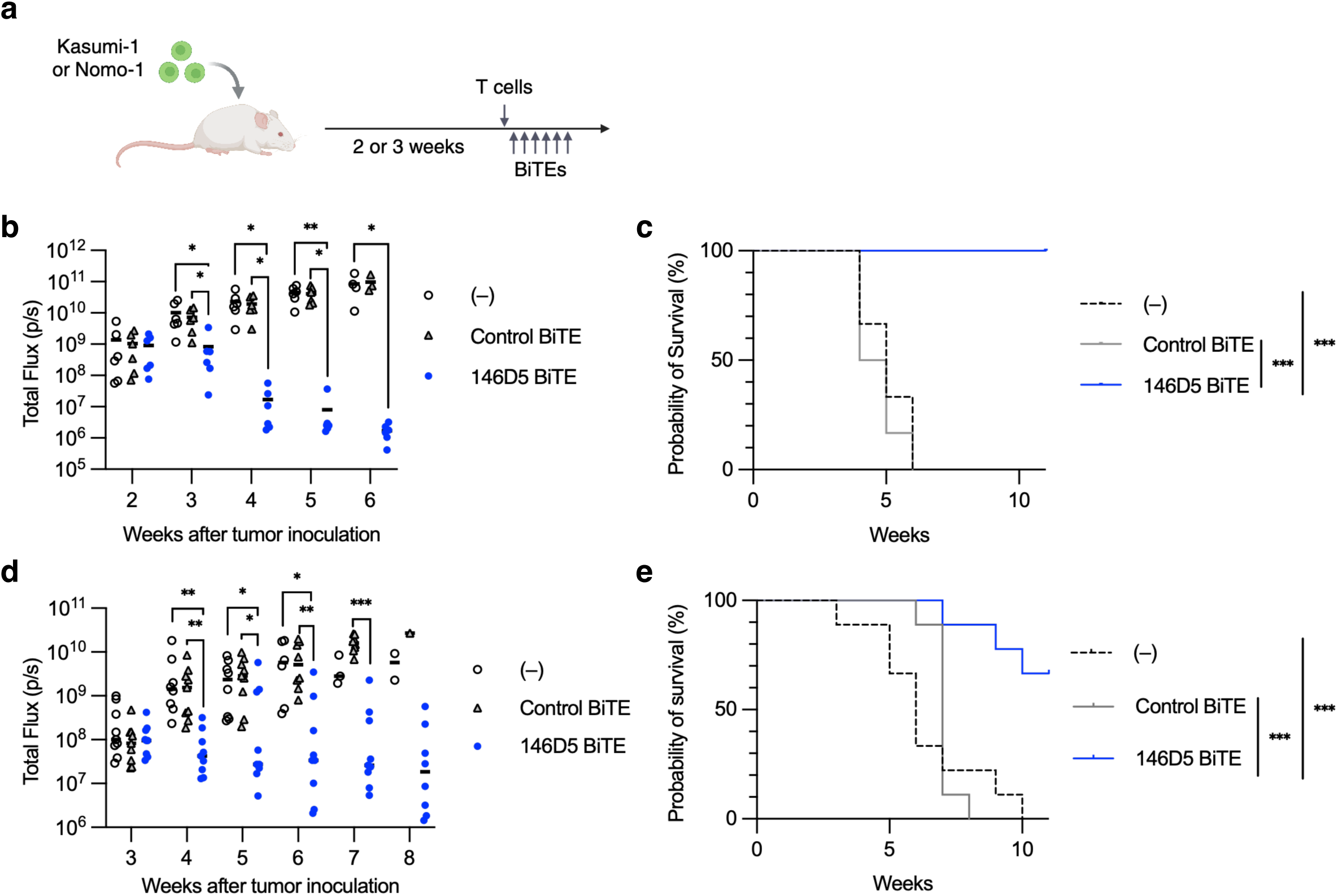
146D5 BiTE shows therapeutic efficacy in AML models **a,** Schematic of NOD-*scid IL2rγnull* (NSG) mice engrafted with Kasumi-1 (a-c) or Nomo-1 cells (a,d,e), infused with T cells after 2 (b,c) or 3 (d,e) weeks, and treated with BiTE for 6 days. **b,c,** Quantification of total photon counts (IVIS imaging of luciferase, b) and Kaplan–Meier survival curve (c) of NSG mice engrafted with Kasumi-1 at indicated time points after tumor inoculation. **d,e,** Quantification of total photon counts (d) and Kaplan–Meier survival curve (e) of NSG mice engrafted with Nomo-1. P-values by one-way ANOVA followed by Dunn’s multiple comparisons (b,d) or log-rank test (c,e): ***p<0.001, **p<0.01, *p<0.05. Each treatment group included 6 (b,c) or 9 (d,e) mice. Representative of 2 independent experiments (b,c) or pooled data from 2 independent experiments (d,e).

## Discussion

T cell-based immunotherapies targeting intracellular antigens through TCR-engineered receptors or conventional TCRm Abs are constrained by strict HLA restriction, limiting their applicability across genetically diverse patient populations. By identifying peptides promiscuously presented by multiple HLA-II alleles and exploiting peptide overhangs extending beyond the HLA-II binding cleft, we developed a novel strategy to overcome this limitation.

We identified a novel MPO-derived epitope, MPO_100-132_, that is promiscuously presented by multiple HLA-II molecules. Each HLA-II allele exhibits unique peptide binding preferences, due to structural variations in their binding clefts. Our competitive binding assays demonstrated that MPO_100-132_ interacts with DP4, DR1, and DR15 through distinct anchor residues, with the peptide containing multiple binding motifs. For instance, hydrophobic or aromatic residues at positions 1 (V113) and 6 (Y118) are expected to act as anchors for common DP alleles such as DP2, DP4, and DP5^39^. The open-ended binding cleft of HLA-II permits longer peptides to be presented with alternative core registers, enabling the same peptide to be presented across multiple HLA-II alleles.

Unlike conventional TCRm Abs, which typically engage both peptide and HLA^11,12^, 146D5 Ab was designed to specifically recognize the N-terminal overhang of the MPO_100-132_ presented by DP4. Mice were immunized with the N- or C-terminal overhang peptides that fall outside the core-binding region on DP4, leading to the successful generation of 146D5 Ab. While the core-binding region of the peptide may adopt distinct conformations across HLA-II alleles, the overhang residues are expected to retain a conserved structure. Based on predicted binding cores and mutagenesis mapping, the 146D5 epitope resides within the N-terminal overhang in most HLA-II alleles, which may explain the observed cross-reactivity of 146D5-based constructs. In some alleles, such as DR1, DR4, and DR16, the epitope overlaps with the predicted peptide core yet is still recognized by 146D5 Ab. Structural modeling of peptide-HLA-II complex for these alleles was inconclusive. However, the antibody epitope may remain exposed despite partial burial within the groove, or that the peptide may adopt multiple binding modes, allowing the epitope to be accessible in certain conformations. This peptide-specific mode of recognition allows 146D5-based therapies to bypass strict HLA restriction and may reduce the risk of off-target toxicity arising from recognition of HLA on healthy tissues.

Despite the phenotypic and genetic heterogeneity of AML, MPO is broadly expressed in blast cells of myelocytic, granulocytic, and monocytic lineage, encompassing French American British (FAB) subtypes M0 to M5^32^. During myeloid differentiation, MPO expression begins at the common myeloid progenitor stage and persists during myelocytic differentiation or up to monocyte committment^31^. Consistent with this expression pattern, 146D5 BiTE mediated cytotoxicity against AML cell lines representing distinct FAB subtypes, including ME-1 (M4), Kasumi-1 (M2), and Nomo-1 (M5)^46,47^, as well as against primary AML samples expressing HLA-II. These findings underscore the broad applicability of 146D5-based approaches.

Importantly, the selective targeting of the peptide, derived from the proMPO, by 146D5 constructs may mitigate off-target toxicity toward healthy myeloid cells. AML cells, but not normal myeloid cells, exhibited detectable levels of immature MPO precursors such as apoproMPO/proMPO, likely due to delayed or inefficient protein maturation. In the ER, apoproMPO undergoes co-translational N-glycosylation, transient interactions with molecular chaperones, and heme incorporation to generate proMPO. ProMPO is subsequently transported to the Golgi, where propeptide is removed by proprotein convertases to generate mature MPO^48^. Impairment at any of these steps may result in the accumulation of immature MPO precursors. Such processing inefficiencies may reflect tumor-specific dysregulation or developmental stage-related immaturity in AML cells. Although 146D5 BiTE recognized a homologous peptide from the EPX propeptide, EPX undergoes a similar protein processing^44^. Furthermore, eosinophils lacked HLA-II expression, reducing the likelihood of unintended targeting in healthy tissue.

Our study highlights the potential of antibody-based therapies targeting peptides presented by HLA-II and suggests that this strategy may be applicable to other tumor types. Compared with the ubiquitous expression of HLA-I, HLA-II expression is more restricted but inducible in various tumor types. This selectivity may lower the risk of toxicity but also limits therapeutic reach. HLA-II expression levels vary depending on tumor origin and molecular subtype^22,49^. In addition to tumors derived from tissues with constitutive HLA-II^50,51^, aberrant HLA-II expression can be induced in other tumor types by cytokines in the inflammatory tumor microenvironment^49^, viral infection^52^, or genetic alterations^53^.

Accordingly, many types of solid tumors have been reported to express HLA-II in situ or in vivo^22,54,55^. Furthermore, HLA-II-peptide targeted strategies could be enhanced by combination therapies. IFN-γ robustly induces HLA-II expression through upregulation of the transcriptional master regulator class II transactivator (CIITA)^50^ and has been shown to increase HLA-II expression in melanoma^56^. Similarly, epigenetic modulator can reverse CIITA silencing and potentiate IFN-γ-mediated HLA-II upregulation in many tumor types^57–59^. Since HLA-II expression correlates with IFN-γ-rich tumor microenvironments^60^, immune checkpoint therapies that augments IFN-γ production^61,62^ may also enhance HLA-II expression, thereby supporting combination approaches.

Further studies employing *in silico* prediction or immunopeptidome screening for promiscuously presented peptides^18^, followed by the generation of cross-recognizing antibodies, may broaden the therapeutic landscape. This approach could yield new treatment options for diverse malignancies, especially when combined with agents that upregulate HLA-II expression.

## Materials and Methods

### Database search

mRNA expression levels of *HLA-DRA* and *MPO* in the Oregon Health & Science University (OHSU) Beat AML cohort^35,36^ were analyzed using cBioPortal for Cancer Genomics. Binding affinity of peptides to HLA-II alleles and core binding sequences were predicted using NetMHCII v2.3 and NetMHCIIpan v4.3^37,38^. Peptides were selected based on predicted affinity (nM) or %Rank. Candidate cross-reactive peptides were identified by BLAST and selected based on query cover and sequence identity. Peptides with high homology to the N-terminal overhang of MPO_100-132_ presented by DP4, or containing 5 residues (Y101, F102, P105, V106, and A107) critical for 146D5 BiTE recognition were selected (See Supplementary Table 6).

### Primary cells and cell lines

Primary leukemia samples, obtained from patient peripheral blood or bone marrow, were provided by Dr. Mark Minden through the Leukemia Tissue Bank at Princess Margaret Cancer Centre following Research Ethics Board (REB) approval. Healthy donor peripheral blood samples were collected under REB approval. Cell lines used in this study included K562, T2, PG13, Jurkat (clone E6-1), Ramos, and Kasumi-1 (all obtained from the American Type Culture Collection); ME-1 and Nomo-1 (from DSMZ, Leibniz Institute); Expi293 (Thermo Fisher Scientific). Jurkat76 cells lacking endogenous TCR, CD4, and CD8 were provided by M. Heemskerk (Leiden University Medical Center)^63^. Jurkat76 cells expressing CD4 (clone 3F7) were generated via retroviral transduction^41^. Engineered cell lines included K562 expressing individual HLA-II alleles (DP, DR, or DQ; see Supplementary Table 1 for allele notations), full-length MPO, or a transmembrane form of anti-146D5 Ab, were generated by retroviral transduction. Artificial antigen-presenting cells (aAPC) were established by transducing CD80 and CD83, as previously described^64^. For T cell expansion, aAPC expressing a transmembrane form of anti-CD3 Ab (aAPC/mOKT3) were used^64^. All cell lines were routinely assessed for mycoplasma by PCR. HLA typing was obtained from Cellosaurus database or conducted by Scisco Genetics Inc. Characterization of primary AML samples determined by flow cytometry is summarized in Supplementary Table 5.

### *In vitro* culture of human T cells and myeloid cells

Peripheral blood mononuclear cells (PBMCs) were isolated by Ficoll-Paque PLUS (GE Healthcare). CD3^+^T cells were isolated using CD3 MicroBeads (Miltenyi Biotec) and stimulated with irradiated with (200 Gy, cesium-137) aAPC/mOKT3 at an E:T of 5:1 in the presence of 100 IU/ml IL-2 and 10 ng/ml IL-15 (PeproTech). Culture media were replenished every 2-3 days. For BiTE cytotoxicity assays, CD8^+^ cells were enriched on day 3-6 using CD8 MicroBeads (Miltenyi Biotec).

Neutrophils were isolated using EasySep Direct Human Neutrophil Isolation Kit (Stemcell Technologies) and stimulated with 5 ng/mL IFN-γ, 55 ng/mL GM-CSF, and 30 ng/mL IL-3 (PeproTech) for 3 days. Monocytes were isolated using CD14 MicroBeads (Miltenyi Biotec) and stimulated with 50 ng/mL M-CSF and 25 ng/mL IL-10 (PeproTech), or 50 ng/mL GM-CSF for 2 to 6 days. T cells and myeloid cells from the same donors were cocultured for assays. Donors carrying at least one HLA-II alleles, capable of binding to MPO_100-132_ with the peptide being recognized by 146D5 constructs in K562 cells systems, were used. All primary cells were maintained in RPMI 1640 with 25mM HEPES (Gibco), supplemented with 10% human AB serum (Gemini Bio-Products) and 50 ng/mL gentamicin (Bioshop).

### Gene transduction and protein production

Genes were cloned into pMX retroviral vector and transduced using 293GPG or PG13 cell-based retrovirus system^41^.

For BiTE or HLA-II production, genes were cloned into pcDNA3.1 or 3.4 vector (Thermo Fisher Scientific) and transfected into Expi293 using ExpiFectamine 293 transfection kit (Thermo Fisher Scientific). Proteins were purified from supernatants using TALON resin (Takara Bio), quantified using NanoDrop (Thermo Fisher Scientific), and analyzed by SDS-PAGE with SimplyBlue SafeStain (Invitrogen). Control BiTEs used scFvs derived from anti-CD19 (HD37) or anti-mesothelin (SS1) fused with anti-CD3 (OKT3). Activity was validated by cytotoxicity assays as shown in Extended Data Fig. 3 in advance. For MPO_100-132_ tethering constructs used in BLI, MPO_100-132_ was linked to the ectodomain of HLA-DP β-chain via a flexible linker (GGGGSAIASGGGG) and an Avi tag at the C-terminus via a GS linker. α-and β-chain were transfected into Expi293 with BirA biotin ligase, and 100 μM biotin were added. Other HLA-II protein for dimer/tetramer staining were produced as previously described^41^.

### Flow cytometry

Fc Receptors were blocked with Human TruStain FcX (BioLegend). Cells were stained with the following antibodies: Alexa Fluor 647-anti HLA-DR,DP,DQ (Tü39, BioLegend), PE anti-HLA-DR,DP,DQ (9-49, Beckman), PE anti-MPO (MPO455-8E6, Thermo Fisher Scientific), FITC anti-CD45 (HI30, BioLegend), Brilliant Violet 421 anti-CD193 (5E8, BioLegend), PE/Cyanine7 anti-Siglec-8 (7C9, BioLegend), PE anti-His (AD1.1.10, Abcam), PE anti-mouse IgG (Poly4053, BioLegend). Dead cells were excluded with 7-AAD (BioLegend) or Fixable Viability dye (Thermo Fisher Scientific). Intracellular staining used BD Cytofix/Cytoperm (BD Biosciences). For eosinophils, red blood cells were lysed with ACK Lysing Buffer (Thermo Fisher Scientific) and SSC^high^ cells were gated. Acquisition was on a CytoFLEX S Flow Cytometer (Beckman Coulter). Sorting used a FACSAria Fusion (BD Biosciences). Data were analyzed using FlowJo (BD Biosciences).

For the analysis of tumor cells isolated from mice, excised tumor tissues were digested using Tumor dissociation kit (Miltenyi Biotec). Mononuclear cells were then isolated by Ficoll-Paque PLUS, incubated with Fc-receptor blocking reagents, and stained with the indicated antibodies.

### Peptides

Synthetic peptides (Genscript) were dissolved in DMSO at 50 mg/ml. Cells were pulsed with peptides (10 µg/ml) for 1-3 hours. The following peptides were used: MPO_109-128_ (TRTAVRAADYLHVALDLLER), MPO_242-261_ (LTPDQERSLMFMQWGQLLDH), MPO_440-459_ (ERLYQEARKIVGAMVQIITY), MPO_494-513_ (FTNAFRYGHTLIQPFMFRLD), MPO_100-132_ (SYFKQPVAATRTAVRAADYLHVALDLLERKLRS), WT1_328-348_ (PGCNKRYFKLSHLQMHSRKHT), CLIP (LPKPPKPVSKMRMATPLLMQALPM). For immunization and ELISA, MPO_100-112_ (SYFKQPVAATRTA) and MPO_122-132_ (ALDLLERKLRS) were BSA-conjugated. MPO_100-132_ alanine-scan variants (residue at 100-112, 113, 114, 118, and 120) were generated, substituting alanine to serine where necessary. C-terminally biotinylated MPO_100-132_ peptides were used for staining with PE or PE/Cyanine7 streptavidin (BioLegend).

### Prediction and validation of T cell epitopes

MPO-derived peptides were predicted using NetMHCII. Frequencies of peptide-specific CD4^+^ T cells in DP4^+^ AML patient samples were measured using DP4 dimers^41^. CD4^+^ T cells were purified using CD4 MicroBeads (Miltenyi Biotec) and stimulated with irradiated DP4 aAPC pulsed with peptides (E:T=20:1). After 48 hours, 10 IU/ml IL-2 and 10 ng/ml IL-15 were added. Cultures were maintained for 2 weeks with medium replenished every 2–3 days. DP4-dimer^+^ T cells were analyzed and sorted for TCR cloning. TCR genes were cloned by 5’-RACE PCR into retrovirus vectors^65^. TCR-transduced Jurkat (3F7) were cocultured with targets and analyzed by IL-2 ELISPOT assays.

### HLA-II dimer and tetramer staining

CD4^+^ T cells were treated with 50 nM dasatinib (LC Laboratories) and stained with affinity maturated DP4 dimers^41^. Purified HLA-II proteins were peptide-exchanged for 18–24 hours and dimerized with PE anti-His Abs. For staining of 146D5-IgG or control (irrelevant peptide-specific)-IgG expressing K562, MPO_100-132_-loaded DP4, DR1, and DR15 monomers were tetramerized with streptavidin.

### ELISPOT assays

IL-2 and IFN-γ ELISPOT assays were performed as previously described^41^. Briefly, PVDF plates (Millipore) were coated with anti-IL-2 (SEL002, R&D Systems) or anti-IFN-γ (1-D1K, MABTECH). T cells were seeded with 2×10^4^ target cells per well and incubated for 20–24 hours at 37°C. Plates were incubated with biotin-anti-IL-2 (SEL002, R&D Systems) or biotin-anti-IFN-γ (7-B6-1, MABTECH) and subsequently HRP-conjugated streptavidin (Jackson ImmunoResearch) was added. Spots were developed with AEC substrate (Sigma-Aldrich) and counted with ImmunoSpot v5.0 (Cellular Technology Limited).

### Generation of antibodies and binding analysis

Antibody generation (mouse immunization, hybridoma generation, and initial screening) was conducted by MEDIMABS. Binding of 146D5 Ab to MPO_100-132_, alanine-scan variants, MPO_100-112_- or MPO_122-132_-BSA conjugates were analyzed by ELISA. Peptides (1 µg/ml) were coated on MaxiSorp 96-well plates (Thermo Fisher Scientific). Plates were incubated with serial dilutions of 146D5 Ab, followed by HRP-anti-mouse IgG (SouthernBiotech). Reactions were developed with 1-step TMB (Thermo Fisher Scientific), terminated by sulfuric acid, and analyzed on a CLARIOstar (BMG Labtech). Background OD was subtracted. Binding of antibodies to pulsed or endogenous MPO/HLA-II was detected with PE anti-mouse IgG and analyzed by flow cytometry. For analysis of BiTE binding, BiTE were dimerized with PE anti-His Ab and used.

### Bio-layer interferometry (BLI)

BLI experiments were performed using an Octet Red96 instrument with streptavidin-coated biosensor tips (Sartorius) at 25 °C using black flat bottom 96-well microtiter plates (Greiner Bio-One) and HBS-EP+ buffer (Cytiva). Sensors were loaded with 50 nM biotinylated ligands (MPO_100-132_ peptide, CLIP_87-101_-tethering-DP4 (NIH), MPO_100-132_-tethering DP4), quenched in 100 nM free biotin (Sigma-Aldrich), and associated with 146D5 Ab at varying concentrations, followed by dissociation in buffer. Data were analyzed using Octet Data Analysis software. Steady-state binding across concentrations was globally fit with a 1:1 Langmuir model.

### Competitive binding assays

K562 or T2 were incubated with 0.5 μM biotinylated-CLIP peptide at 37 °C for 2 hours, either alone or together with competitive peptides at the indicated concentrations. After washing, cells were stained with PE or PE/Cyanine7-conjugated streptavidin and analyzed by flow cytometry. Dose-response inhibition curves were fitted by nonlinear regression, and LogIC_50_ values were derived from the fitted models. Percentage inhibition was calculated from MFI as: % Inhibition =100 x [1-(MFI with competitive peptide / MFI with CLIP peptide only)].

### Structural prediction

Interactions between MPO_100-132_ and HLA-II molecules was predicted using AlphaFold 3 installed on a local cluster^43^. Predictions used 1,000 or more seeds with one diffusion sample per seed. Top-ranked models were selected based on the values of ranking_score, iptm, chain_pair_iptm, and pLDDT. Models were visualized using UCSF ChimeraX-1.9^66^.

### *In vitro* cytotoxicity assays

Target cells were labeled with 5 μM Vybrant DiO cell-labeling solution (Thermo Fisher Scientific). Effector T cells were added at the indicated E:T ratio with BiTE at the specified concentrations. After 16-18 hours of coculture, cells were stained with 3 μM TO-PRO-3 (Thermo Fisher Scientific) and analyzed by flow cytometry. Dead target cells were defined as the percentage of TO-PRO-3^+^ cells among DiO^+^ cells. Cytotoxicity was calculated as: (% dead target cells in test wells) – (% dead target cells without effector cells). For assays using primary AML samples or normal myeloid cells, cytotoxicity was defined as: (% dead target cells in the wells with BiTE) – (% dead target cells in the wells without BiTE), using autologous T cells. In primary AML cell assays, anti-SS1 BiTE were used as negative controls to account for the presence of CD19^+^ AML cells. In other experiments, anti-CD19 BiTE were used as negative controls.

### Immunoblotting

Cells were lysed in RIPA buffer (Thermo Fisher Scientific) supplemented with protease inhibitors (Bioshop). Lysates were mixed with 4x Laemmli sample buffer (Bio-Rad) and 2-mercaptoethanol (Sigma-Aldrich) and boiled. Proteins were resolved by SDS-PAGE, transferred to PVDF membranes (Millipore), and probed with primary Abs: anti-MPO (EPR4793, Abcam) or anti-GAPDH (D16H11, CST). HRP-conjugated anti-rabbit IgG (Promega) was used as secondary Ab, and signals were detected with Clarity Western ECL substrate (Bio-Rad) and imaged using ChemiDoc system (Bio-Rad).

### Mouse experiments

Female NSG mice bred at the Princess Margaret Cancer Centre Animal Resource Centre (ARC) were used. Mice were subcutaneously injected with 5×10^6^ Kasumi-1 or 2×10^6^ Nomo-1 transduced with GFP-firefly luciferase. Two weeks (Kasumi-1) or three weeks (Nomo-1) later, mice intravenously received 1×10^7^ T cells, stimulated with aAPC/mOKT3. *In vivo* tumor growth was monitored by bioluminescent imaging using the IVIS Spectrum (Perkin Elmer) after administration of D-Luciferin (Gold Biotechnology), and data were analyzed using Living Image software (Perkin Elmer). Tumor volumes were monitored every 2–3 days until it exceeded 15 mm. Mice were monitored daily and euthanized by CO_2_ inhalation if humane endpoints were reached (tumor progression, bilateral hind-limb paralysis, or >20% weight loss). Treatment groups were assigned based on tumor size the day before T cell injection. No statistical methods were used to predetermine sample size. Investigators were not blinded to treatment allocation or outcome assessment.

### Statistics

All experiments were conducted using T cells from at least two different donors. Statistical analyses (one-way ANOVA, two-way ANOVA, nonlinear regression, linear regression, and Kaplan-Meier survival analysis) were performed using GraphPad Prism 10. Differences were considered statistically significant at P < 0.05. Sample sizes were not predetermined statistically.

### Study approval

All studies were performed in accordance with the Helsinki Declaration and approved by the Research Ethics Board of the University Health Network (UHN), Toronto, Canada. Written informed consent was obtained from all healthy donors. All animal experiments were approved by the Princess Margaret Cancer Centre Animal Care Committee, UHN and performed according to Canadian Council on Animal Care guidelines.

### Author contributions

SF, EYFZ, and NH designed the research. SF, EYFZ, FI, YM, YO, DHH, XW, KH, BDB, DL, and YK performed the experiments. All the authors analyzed the results. SF, DL, and NH designed the figures and drafted the manuscript. All the authors reviewed and contributed to the manuscript.

## Acknowledgments

This work was supported by the Canadian Institutes of Health Research Project Grant PJT190190 (to N.H), the Ontario Institute for Cancer Research Clinical Investigator Award IA-039 (to N.H.), the Longo Family Cancer Foundation (to N.H.), the Ira Schneider Memorial Cancer Research Foundation (to N.H.), the Princess Margaret Cancer Foundation (to N.H.), the Longo Family Cancer Immunotherapy Fellowship (to Y.M.). We acknowledge the Flow Cytometry Core Facility at the Princess Margaret Cancer Centre and Princess Margaret Cancer Centre ARC for their assistance with this study. Schematics were created with BioRender.com.

---

**Extended Data Fig. 1.**
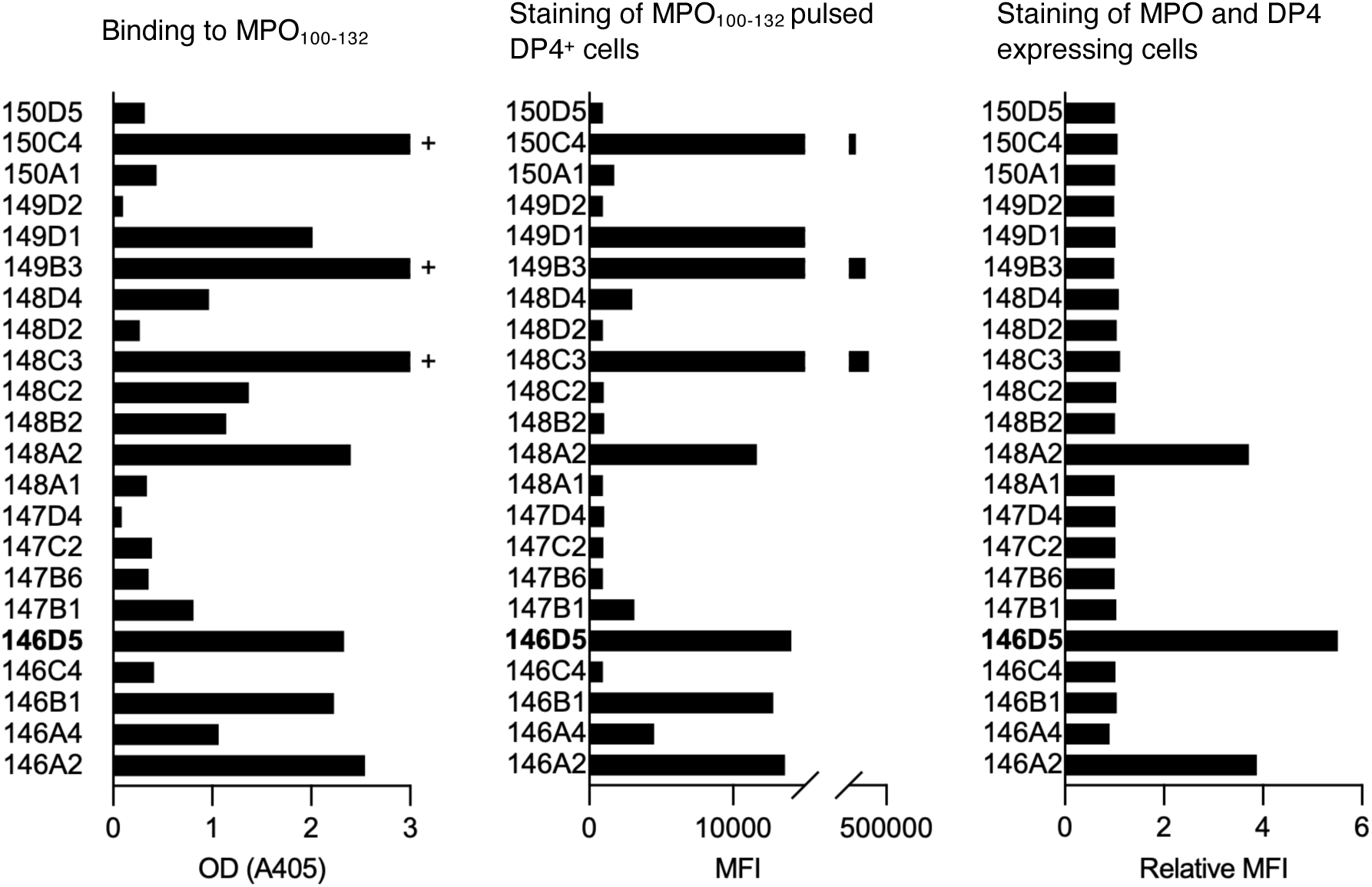
Validation of antibodies generated against MPO_100-132_ Binding of the antibodies from hybridoma supernatants to plate-coated MPO_100-132_ (left), DP4^+^ K562 cells pulsed with MPO_100-132_ (middle), and K562 cells expressing both DP4 and MPO (right). ELISA OD values (left), and flow cytometry MFI (middle, right) are shown. Relative MFI was calculated as the ratio of MFI from DP4^+^ MPO^+^ K562 to DP4^+^ K562.

**Extended Data Fig. 2.**
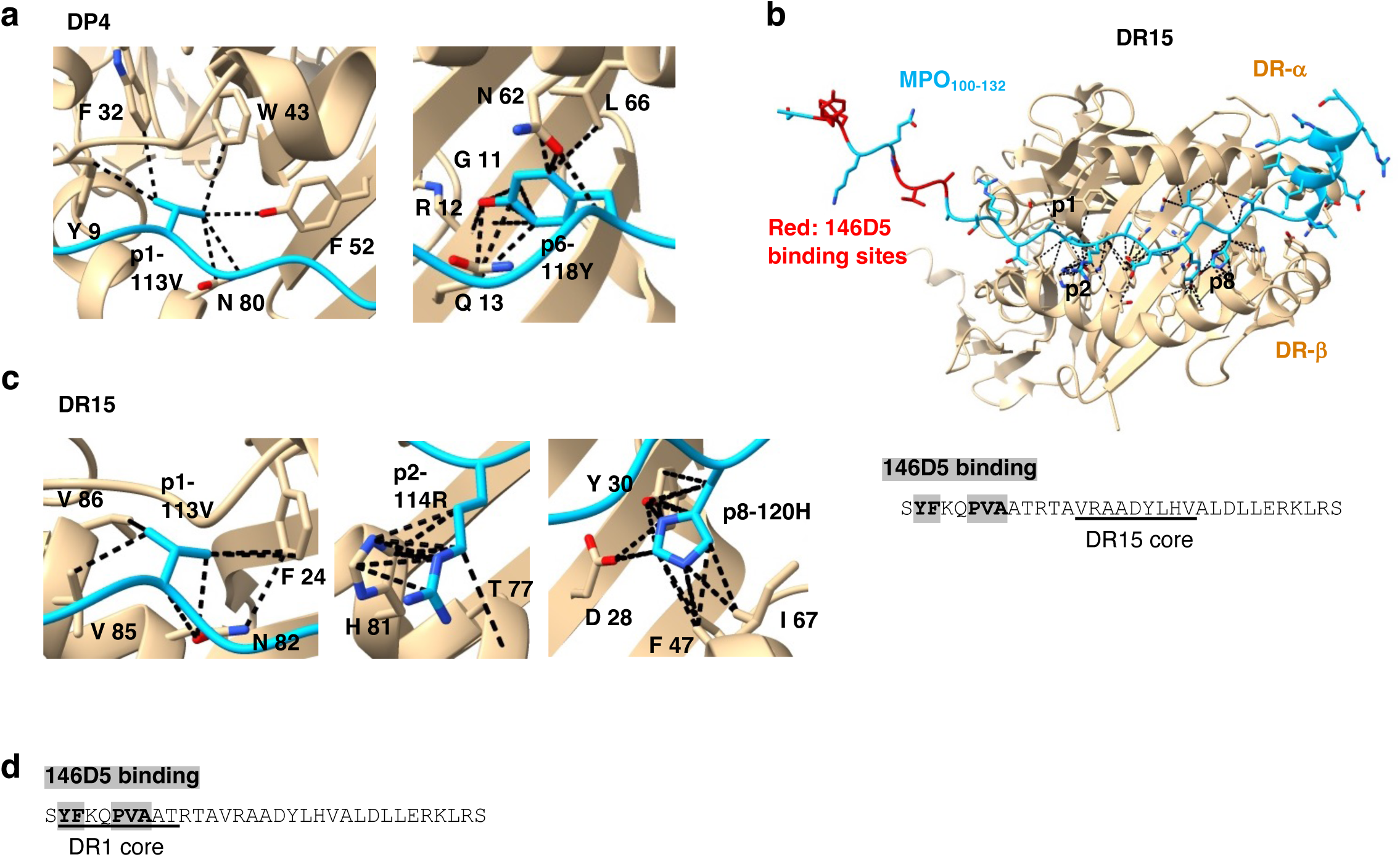
Structural modeling of MPO_100-132_ with HLA-II **a,** Structural model of MPO_100-132_ bound to DP4 generated using AlphaFold. Close-up views of peptide in positions 1 (p1, 113V, left) and p6 (P118Y, right) binding pocket. **b,c,** Structural models of MPO_100-132_ bound to DR15 generated by AlphaFold. Overall structure (b) and close-up views of peptide in p1 (113V, left), p2 (114R, middle), and p8 (120H, right) binding pocket (c). The peptide is shown in cyan with 146D5 Ab-contact residues in red; HLA-II in light brown. Black dashes represent contacts defined as VDW overlap ≥ –0.4 Å using UCSF Chimera X, **b,d,** Sequence of MPO_100-132_ with the predicted binding core (underlined) of DR15 (b, bottom) and DR1 (d). Residues critical for 146D5 Ab binding (shaded gray).

**Extended Data Fig. 3.**
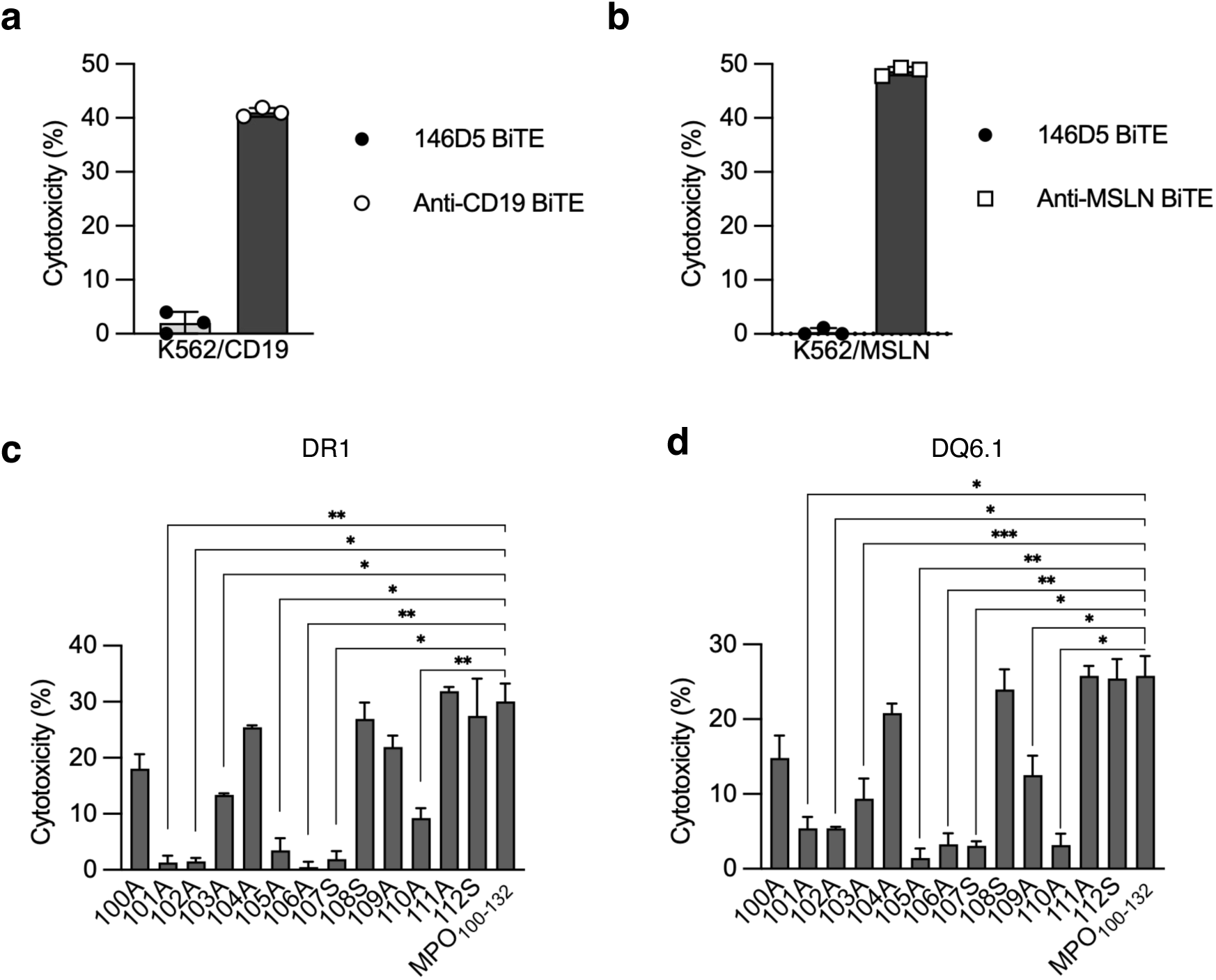
Epitope mapping of 146D5 BiTEs **a,b,** Cytotoxicity of K562 cells expressing CD19 (a), mesothelin (MSLN, b) by co-cultured T cells in the presence of 1 nM indicated BiTE. **c,d,** Cytotoxicity of K562 cells expressing DR1^+^ or DQ6.1^+^K562 cells pulsed with MPO_100-132_ mutants by co-cultured T cells in the presence of 1 nM 146D5 BiTE. Cytotoxicity was shown. All cytotoxicity assays at E:T=2:1. Data represent mean ± S.D. from 3 donors. P-values by one-way ANOVA followed by Dunnett’s multiple comparisons (c,d): ***p<0.001, **p<0.01, *p<0.05. Representative data of 2 independent experiments.

**Extended Data Fig. 4.**
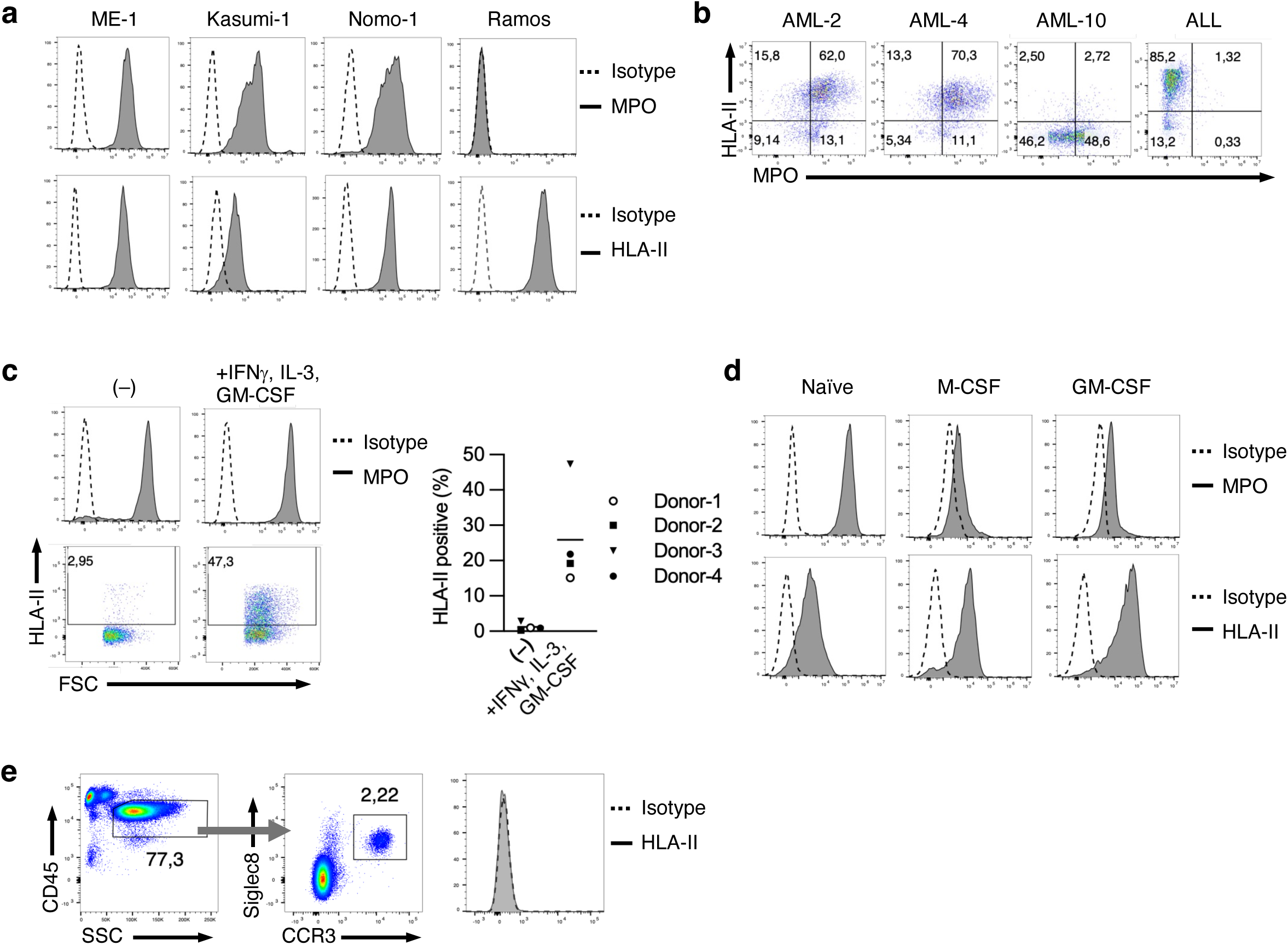
Characterization of AML samples and primary cells **a,b,** Representative staining of MPO and HLA-II in indicated cell lines, or primary AML or ALL samples. **c,d,** Representative staining of MPO and HLA-II in naïve or activated neutrophils (c, left) and monocytes (d) from healthy donors. Frequencies of HLA-II^+^ cells are shown (c, right). **e,** Representative staining of HLA-II in eosinophils (CD45^+^ SSC^high^ Siglec8^+^ CCR3^+^) from healthy donor blood. Representative of at least 2 independent experiments (a-e).

**Extended Data Fig. 5.**
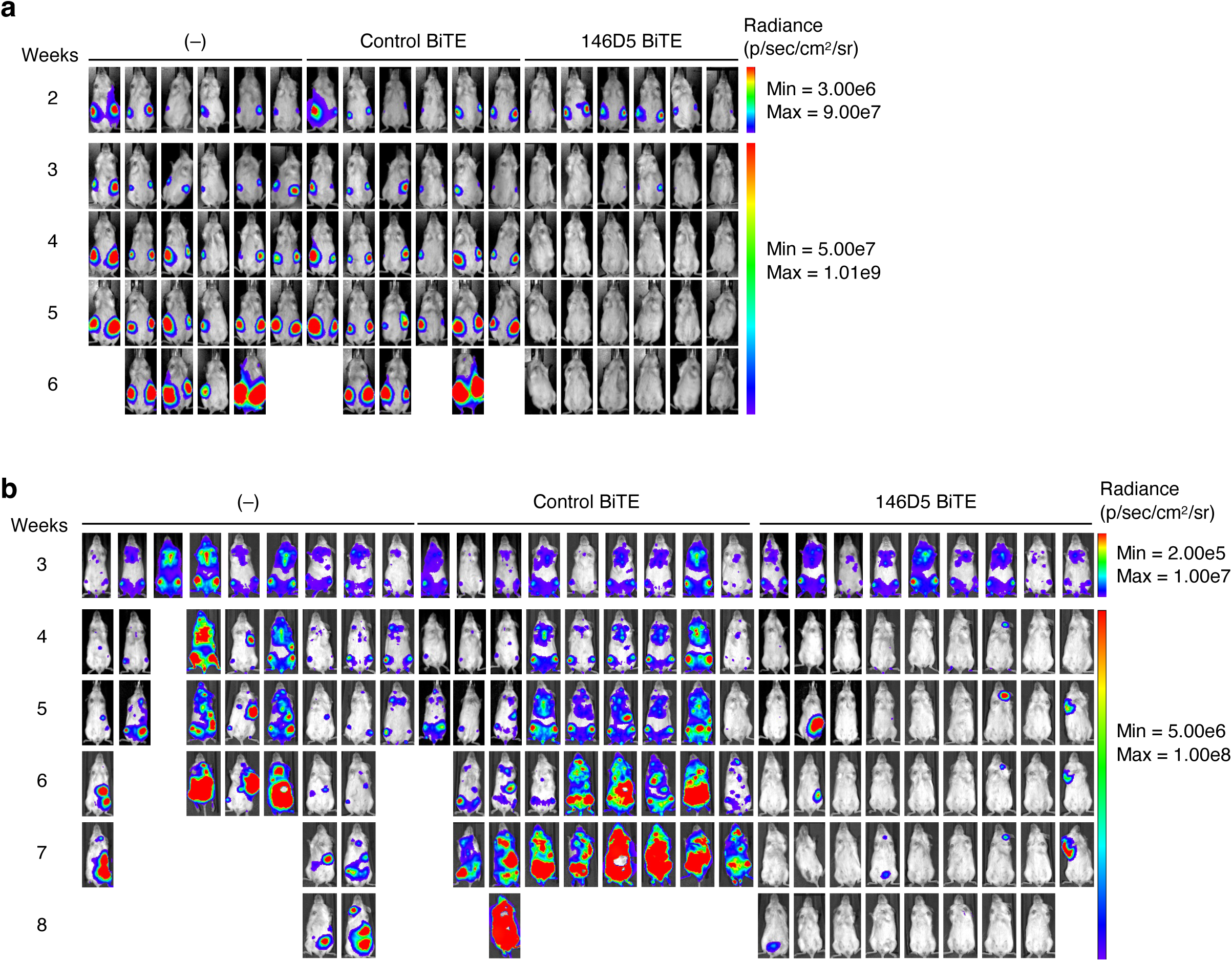
Therapeutic efficacy of 146D5 BiTE in AML models **a,b,** Bioluminescence images showing tumor burden at the indicated weeks after injection of Kasumi-1 (a) or Nomo-1 (b) cells.

**Supplementary Table. 1.** Notation of HLA alleles in this study.

| Name | $\alpha$ chain | $\beta$ chain |
| --- | --- | --- |
| DP2 | DPA1*01:03 | DPB1*02:01 |
| DP4 | DPA1*01:03 | DPB1*04:01 |
| DP5 | DPA1*01:03 | DPB1*05:01 |
| DR1 | DRA1*01:01 | DRB1*01:01 |
| DR3 | DRA1*01:01 | DRB1*03:01 |
| DR4 | DRA1*01:01 | DRB1*05:01 |
| DR7 | DRA1*01:01 | DRB1*07:01 |
| DR11 | DRA1*01:01 | DRB1*11:01 |
| DR13 | DRA1*01:01 | DRB1*13:01 |
| DR15 | DRA1*01:01 | DRB1*15:01 |
| DR16 | DRA1*01:01 | DRB1*16:01 |
| DQ4 | DQA1*0401 | DQB1*0402 |
| DQ5 | DQA1*0101 | DQB1*0501 |
| DQ6 | DQA1*0103 | DQB1*0601 |
| DQ9 | DQA1*02:01 | DQB1*03:03 |

**Supplementary Table. 2.** Predicted binding affinities of MPO-derived peptides to HLA-II alleles by NetMHCII.

|  | DP4 | DP5 | DR1 | DR15 |
| --- | --- | --- | --- | --- |
| **MPO<sub>109-128</sub>** | **56.2** | **73.8** | **197.1** | **253.8** |
| MPO<sub>242-261</sub> | 41.8 | 166.6 | 138.5 | 412.6 |
| MPO<sub>440-459</sub> | 104.1 | 80.5 | 192.9 | 450.6 |
| MPO<sub>494-513</sub> | 69.9 | 41.6 | 220.1 | 64 |

**Supplementary Table. 3.** Predicted binding affinities and core binding sequences of MPO_100-132_ to HLA-II alleles by NetMHCII.

| Allele | Core | Affinity (nM) |
| --- | --- | --- |
| DP2 | YLHVALDLL | 91.7 |
| **DP4** | **VRAADYLHV** | **38.2** |
| DP5 | HVALDLLER | 18 |
| DR1 | YFKQPVAAT | 22.7 |
| DR3 | VALDLLERK | 21.3 |
| DR4 | YFKQPVAAT | 106.9 |
| DR7 | VRAADYLHV | 122.5 |
| DR11 | LDLLERKLR | 64.3 |
| DR13 | LDLLERKLR | 525.2 |
| DR15 | VRAADYLHV | 112 |
| DR16 | FKQPVAATR | 45.4 |
| DQ4.2 | ADYLHVALD | 438.8 |
| DQ5.1 | DYLHVALDL | 76.3 |
| DQ6.1 | ATRTAVRAA | 21.8 |
| DQ9.2 | AATRTAVRA | 45.2 |

**Supplementary Table. 4.** HLA-II genotyping of cell lines.

| Name | *DPA1* | *DPB1* | *DQA1* | *DQB1* | *DRB1* |
| --- | --- | --- | --- | --- | --- |
| ME-1 | 01:03:01,<br>02:02:02 | 05:01:01,<br>02:02 | 01:03 | 06:02:01 | 08:02:01,<br>15:01:01 |
| Kasumi-1 | 01:03:01 | 02:01:02 | 01:04:01,<br>03:02:01 | 03:03:02,<br>05:03:01 | 09:01:02,<br>14:54:01 |
| Nomo-1 | 01:03:01,<br>02:02:02 | 02:01:02,<br>05:01:01 | 03:03:01 | 04:01:01 | 04:05:01 |
| Ramos | 01:03:01 | 04:01:01,<br>104:01 | 02:01:01 | 02:02:01 | 07:01:01 |

**Supplementary Table. 5.** HLA-II genotyping and characterization of primary AML samples.

|  | MPO<br>(MFI) | HLA-II<br>(MFI) | MPO<sup>+</sup><br>HLA-II<sup>+</sup> (%) | DPA1 | DPB1 | DQA1 | DQB1 | DRB1 |
| --- | --- | --- | --- | --- | --- | --- | --- | --- |
| AML-1 | 8434 | 35059 | 85.2 | 01:03:01,<br>02:02:02 | 05:01:01 | 01:03:01,<br>03:02:01 | 03:03:02,<br>06:01:01 | 08:03:02,<br>09:01:02 |
| AML-2 | 10756 | 24065 | 86.4 | 01:03:01 | 03:01:01,<br>06:01:01 | 03:01:01,<br>05:05:01 | 03:01:01,<br>03:02:01 | 04:04:01,<br>11:01:01 |
| AML-3 | 11341 | 14928 | 86.3 | 02:02:02 | 01:01:01,<br>35:01:01 | 01:02:01,<br>06:01:01 | 03:01:01,<br>05:02:01 | 12:02:01,<br>15:02:01 |
| AML-4 | 8393 | 12745 | 85 | 02:01:01,<br>02:02:02 | 01:01:01,<br>09:01:01 | 01:01:01 | 05:01:01,<br>05:01:24 | 01:01:01,<br>15:02:01 |
| AML-5 | 7183 | 9597 | 72.7 | 01:03:01,<br>02:01:01 | 04:02:01,<br>09:01:01 | 01:02:01,<br>01:03:01 | 06:01:01,<br>06:02:01 | 15:01:01,<br>15:02:01 |
| AML-6 | 9188 | 7607 | 70.7 | 01:03:01 | 02:01:02,<br>04:01:01 | 01:02:01,<br>01:02:02 | 05:02:01,<br>06:02:01 | 15:01:01,<br>16:01:01 |
| AML-7 | 5799 | 6061 | 66.3 | 01:03:01 | 04:01:01,<br>04:02:01 | 02:01:01,<br>05:05:01 | 03:01:01,<br>03:03:02 | 07:01:01,<br>13:03:01 |
| AML-8 | 14019 | 5626 | 64.1 | 01:03:01 | 04:01:01,<br>04:02:01 | 01:02:01,<br>05:05:01 | 03:01:01,<br>05:04 | 01:01:01,<br>11:03:01 |
| AML-9 | 5868 | 4226 | 70.1 | 01:03:01,<br>02:01:01 | 04:01:01,<br>05:01:01 | 01:02:01,<br>05:05:01 | 03:01:01,<br>06:09:01 | 11:04:01,<br>13:02:01 |
| AML-10 | 61152 | 521 | 2.3 | 02:02:02 | 01:01:01,<br>05:01:01 | 01:02:01,<br>06:01:01 | 03:01:01,<br>05:02:01 | 12:02:01,<br>15:02:01 |
| AML-11 | 7782 | 160 | 1.54 | 01:03:01,<br>02:02:02 | 01:01:01,<br>03:01:01 | 02:01:01,<br>05:05:01 | 02:02:01,<br>03:01:01 | 07:01:01,<br>11:01:01 |

**Supplementary Table. 6.**
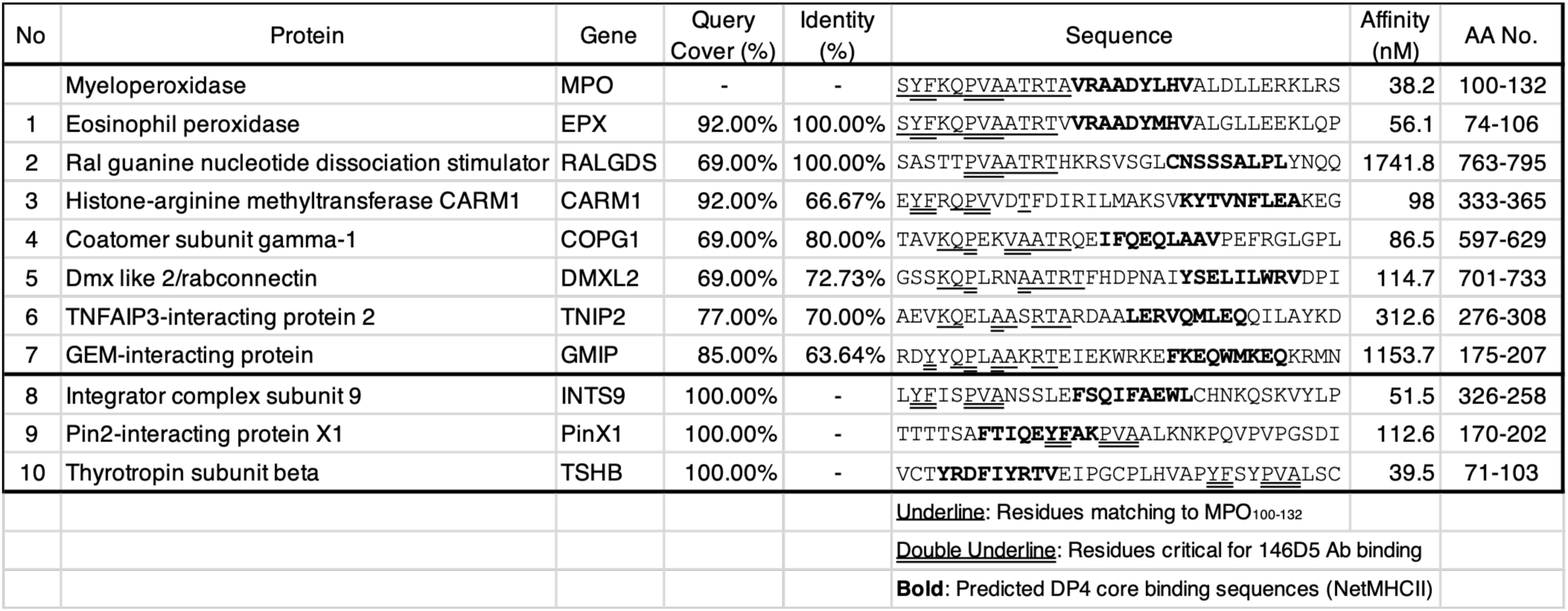
Peptides with high sequence homology to MPO_100-132_.

## References

1. Weiner, G. J. Building better monoclonal antibody-based therapeutics. Nat. Rev. Cancer 15, 361–370 (2015).

2. Paul, S. et al. Cancer therapy with antibodies. Nat. Rev. Cancer 24, 399–426 (2024).

3. van de Donk, N. W. C. J. & Zweegman, S. T-cell-engaging bispecific antibodies in cancer. The Lancet 402, 142–158 (2023).

4. Baeuerle, P. A., Sauer, K., Grieshaber-Bouyer, R. & Michaelson, J. S. T cell engagers emerge as a compelling therapeutic modality. Journal of Experimental Medicine 223, (2026).

5. Goebeler, M.-E. & Bargou, R. C. T cell-engaging therapies — BiTEs and beyond. Nat. Rev. Clin. Oncol. 17, 418–434 (2020).

6. Posey, A. D., Young, R. M. & June, C. H. Future perspectives on engineered T cells for cancer. Trends Cancer 10, 687–695 (2024).

7. Fenis, A., Demaria, O., Gauthier, L., Vivier, E. & Narni-Mancinelli, E. New immune cell engagers for cancer immunotherapy. Nat. Rev. Immunol. 24, 471–486 (2024).

8. Molldrem, J. & Zha, D. Unlocking Intracellular Oncology Targets: The Unique Role of Antibody-Based T-Cell Receptor Mimic (TCRm) Therapeutics in T-Cell Engagers (TCEs) and Antibody-Drug Conjugates (ADCs). Cancers (Basel). 16, 3776 (2024).

9. Akatsuka, Y. TCR-Like CAR-T Cells Targeting MHC-Bound Minor Histocompatibility Antigens. Front. Immunol. 11, (2020).

10. Blum, J. S., Wearsch, P. A. & Cresswell, P. Pathways of Antigen Processing. Annu. Rev. Immunol. 31, 443–473 (2013).

11. Mareeva, T., Martinez-Hackert, E. & Sykulev, Y. How a T cell receptor-like antibody recognizes major histocompatibility complex-bound peptide. J. Biol. Chem. 283, 29053–9 (2008).

12. Ataie, N. et al. Structure of a TCR-Mimic Antibody with Target Predicts Pharmacogenetics. J. Mol. Biol. 428, 194–205 (2016).

13. Yarmarkovich, M. et al. Targeting of intracellular oncoproteins with peptide-centric CARs. Nature 623, 820–827 (2023).

14. Sun, Y. et al. Structural principles of peptide-centric chimeric antigen receptor recognition guide therapeutic expansion. Sci. Immunol. 8, eadj5792 (2023).

15. Mori, S. et al. Neoself-antigens are the primary target for autoreactive T cells in human lupus. Cell 187, 6071–6087.e20 (2024).

16. Racle, J. et al. Machine learning predictions of MHC-II specificities reveal alternative binding mode of class II epitopes. Immunity 56, 1359–1375.e13 (2023).

17. Omiya, R., Buteau, C., Kobayashi, H., Paya, C. V & Celis, E. Inhibition of EBV-Induced Lymphoproliferation by CD4+ T Cells Specific for an MHC Class II Promiscuous Epitope. The Journal of Immunology 169, 2172–2179 (2002).

18. Laghmouchi, A. et al. Promiscuity of Peptides Presented in HLA-DP Molecules from Different Immunogenicity Groups Is Associated With T-Cell Cross-Reactivity. Front. Immunol. 13, (2022).

19. Sette, A., Southwood, S., Miller, J. & Appella, E. Binding of major histocompatibility complex class II to the invariant chain-derived peptide, CLIP, is regulated by allelic polymorphism in class II. J. Exp. Med. 181, 677–683 (1995).

20. Wang, N., Waghray, D., Caveney, N. A., Jude, K. M. & Garcia, K. C. Structural insights into human MHC-II association with invariant chain. Proceedings of the National Academy of Sciences 121, (2024).

21. Ghosh, P., Amaya, M., Mellins, E. & Wiley, D. C. The structure of an intermediate in class II MHC maturation: CLIP bound to HLA-DR3. Nature 378, 457–462 (1995).

22. Axelrod, M. L., Cook, R. S., Johnson, D. B. & Balko, J. M. Biological Consequences of MHC-II Expression by Tumor Cells in Cancer. Clinical Cancer Research 25, 2392–2402 (2019).

23. Wen, M., Li, Y., Qin, X., Qin, B. & Wang, Q. Insight into Cancer Immunity: MHCs, Immune Cells and Commensal Microbiota. Cells 12, 1882 (2023).

24. Crotzer, V. L. & Blum, J. S. Autophagy and adaptive immunity. Immunology 131, 9–17 (2010).

25. Blum, J. S. & Zhou, D. Presentation of Cytosolic Antigens Via MHC Class II Molecules. Immunologic Research vol. 30 279–290 https://link.springer.com/10.1385/IR:30:3:279 (2004).

26. Lich, J. D., Elliott, J. F. & Blum, J. S. Cytoplasmic processing is a prerequisite for presentation of an endogenous antigen by major histocompatibility complex class II proteins. J. Exp. Med. 191, 1513–24 (2000).

27. Yamashita, Y. et al. HLA-DP84Gly constitutively presents endogenous peptides generated by the class I antigen processing pathway. Nat. Commun. 8, 1–14 (2017).

28. Hunder, N. N. et al. Treatment of Metastatic Melanoma with Autologous CD4+ T Cells against NY-ESO-1. New England Journal of Medicine 358, 2698–2703 (2008).

29. Lu, Y.-C. et al. Treatment of Patients With Metastatic Cancer Using a Major Histocompatibility Complex Class II-Restricted T-Cell Receptor Targeting the Cancer Germline Antigen MAGE-A3. J. Clin. Oncol. 35, 3322–3329 (2017).

30. Döhner, H. et al. Diagnosis and management of AML in adults: 2017 ELN recommendations from an international expert panel. Blood 129, 424–447 (2017).

31. Vergez, F. et al. Phenotypically-defined stages of leukemia arrest predict main driver mutations subgroups, and outcome in acute myeloid leukemia. Blood Cancer J. 12, 117 (2022).

32. Krause, D. S. & Van Etten, R. A. Right on target: eradicating leukemic stem cells. Trends Mol. Med. 13, 470–481 (2007).

33. Lu, S. et al. Novel myeloperoxidase-derived HLA-A2-restricted peptides as therapeutic targets against myeloid leukemia. Cytotherapy 23, 793–798 (2021).

34. Hanson, C. A., Gajl-Peczalska, K. J., Parkin, J. L. & Brunning, R. D. Immunophenotyping of acute myeloid leukemia using monoclonal antibodies and the alkaline phosphatase-antialkaline phosphatase technique. Blood 70, 83–9 (1987).

35. Tyner, J. W. et al. Functional genomic landscape of acute myeloid leukaemia. Nature 562, 526–531 (2018).

36. Bottomly, D. et al. Integrative analysis of drug response and clinical outcome in acute myeloid leukemia. Cancer Cell 40, 850–864.e9 (2022).

37. Jensen, K. K. et al. Improved methods for predicting peptide binding affinity to MHC class II molecules. Immunology 154, 394–406 (2018).

38. Reynisson, B., Alvarez, B., Paul, S., Peters, B. & Nielsen, M. NetMHCpan-4.1 and NetMHCIIpan-4.0: improved predictions of MHC antigen presentation by concurrent motif deconvolution and integration of MS MHC eluted ligand data. Nucleic Acids Res. 48, W449–W454 (2020).

39. Sidney, J. et al. Five HLA-DP Molecules Frequently Expressed in the Worldwide Human Population Share a Common HLA Supertypic Binding Specificity. The Journal of Immunology 184, 2492–2503 (2010).

40. Castelli, F. A. et al. HLA-DP4, the Most Frequent HLA II Molecule, Defines a New Supertype of Peptide-Binding Specificity. The Journal of Immunology 169, 6928–6934 (2002).

41. Sugata, K. et al. Affinity-matured HLA class II dimers for robust staining of antigen-specific CD4+ T cells. Nat. Biotechnol. 39, 958–967 (2021).

42. Hsiue, E. H.-C. et al. Targeting a neoantigen derived from a common TP53 mutation. Science 371, (2021).

43. Abramson, J. et al. Accurate structure prediction of biomolecular interactions with AlphaFold 3. Nature 630, 493–500 (2024).

44. Nauseef, W. M. Biosynthesis of human myeloperoxidase. Arch. Biochem. Biophys. 642, 1–9 (2018).

45. Davies, M. J., Hawkins, C. L., Pattison, D. I. & Rees, M. D. Mammalian Heme Peroxidases: From Molecular Mechanisms to Health Implications. Antioxid. Redox Signal. 10, 1199–1234 (2008).

46. Skopek, R. et al. Choosing the Right Cell Line for Acute Myeloid Leukemia (AML) Research. Int. J. Mol. Sci. 24, 5377 (2023).

47. Georges, E. et al. Transcriptomic characterisation of acute myeloid leukemia cell lines bearing the same t(9;11) driver mutation reveals different molecular signatures. BMC Genomics 26, 300 (2025).

48. Hansson, M., Olsson, I. & Nauseef, W. M. Biosynthesis, processing, and sorting of human myeloperoxidase. Arch. Biochem. Biophys. 445, 214–224 (2006).

49. Seliger, B., Kloor, M. & Ferrone, S. HLA class II antigen-processing pathway in tumors: Molecular defects and clinical relevance. Oncoimmunology 6, e1171447 (2017).

50. Ting, J. P.-Y. & Trowsdale, J. Genetic Control of MHC Class II Expression. Cell 109, S21–S33 (2002).

51. Wosen, J. E., Mukhopadhyay, D., Macaubas, C. & Mellins, E. D. Epithelial MHC Class II Expression and Its Role in Antigen Presentation in the Gastrointestinal and Respiratory Tracts. Front. Immunol. 9, (2018).

52. Näsman, A. et al. HLA Class I and II Expression in Oropharyngeal Squamous Cell Carcinoma in Relation to Tumor HPV Status and Clinical Outcome. PLoS One 8, e77025 (2013).

53. Reuss, D. E. et al. Functional MHC Class II Is Upregulated in Neurofibromin-Deficient Schwann Cells. Journal of Investigative Dermatology 133, 1372–1375 (2013).

54. van Vreeswijk, Hans., Ruiter, D. J., Bröcker, E.-B., Welvaart, Kees. & Ferrone, Soldano. Differential Expression of HLA-DR, DQ, and DP Antigens in Primary and Metastatic Melanoma. Journal of Investigative Dermatology 90, 755–760 (1988).

55. Younger, A. R. et al. HLA class II antigen presentation by prostate cancer cells. Prostate Cancer Prostatic Dis. 11, 334–341 (2008).

56. Propper, D. J. et al. Low-dose IFN-gamma induces tumor MHC expression in metastatic malignant melanoma. Clin. Cancer Res. 9, 84–92 (2003).

57. Pickles, O. J. et al. MHC Class II is Induced by IFNγ and Follows Three Distinct Patterns of Expression in Colorectal Cancer Organoids. Cancer Research Communications 3, 1501–1513 (2023).

58. Liu, J. H., Bian, Y. M., Xie, Y. & Lu, D. P. Epigenetic modification and preliminary investigation of the mechanism of the immune evasion of HL-60 cells. Mol. Med. Rep. 12, 1059–1065 (2015).

59. Tovar Perez, J. E., et al. Epigenetic regulation of major histocompatibility complexes in gastrointestinal malignancies and the potential for clinical interception. Clin. Epigenetics 16, 83 (2024).

60. Galon, J. et al. Type, Density, and Location of Immune Cells Within Human Colorectal Tumors Predict Clinical Outcome. Science (1979). 313, 1960–1964 (2006).

61. Grasso, C. S. et al. Conserved Interferon-γ Signaling Drives Clinical Response to Immune Checkpoint Blockade Therapy in Melanoma. Cancer Cell 38, 500–515.e3 (2020).

62. Wong, C. W., Huang, Y. Y. & Hurlstone, A. The role of IFN-γ-signalling in response to immune checkpoint blockade therapy. Essays Biochem. 67, 991–1002 (2023).

63. Heemskerk, M. H. M. et al. Redirection of antileukemic reactivity of peripheral T lymphocytes using gene transfer of minor histocompatibility antigen HA-2-specific T-cell receptor complexes expressing a conserved alpha joining region. Blood 102, 3530–3540 (2003).

64. Butler, M. O. et al. Ex vivo expansion of human CD8 + T cells using autologous CD4 + T cell help. PLoS One 7, (2012).

65. Murata, K. et al. Landscape mapping of shared antigenic epitopes and their cognate TCRs of tumor-infiltrating T lymphocytes in Melanoma. Elife 9, 1–22 (2020).

66. Meng, E. C. et al. UCSF ChimeraX: Tools for structure building and analysis. Protein Science 32, (2023).

